# A MAD7-based genome editing system for *E. coli*

**DOI:** 10.1101/2022.09.01.505100

**Authors:** Markus Mund, Wadim Weber, Daniel Degreif, Christoph Schiklenk

## Abstract

A broad variety of biomolecules is industrially produced in bacteria and yeasts. These microbial expression hosts can be optimized through genomic engineering using CRISPR tools. Here, we designed and characterized such a modular genome editing system based on the Cas12a-like RNA guided nuclease MAD7 in *E. coli*. Our system enables the efficient generation of single nucleotide polymorphisms (SNPs) or gene deletions and can directly be used with donor DNA from benchtop DNA assembly to increase throughput. We combined multiple edits to engineer an *E. coli* strain with reduced overflow metabolism and increased plasmid yield, highlighting the versatility and industrial applicability of our approach.

## Introduction

Microbial cell factories have enormous economical relevance for the chemical, biotechnological and pharmaceutical industry and will be pivotal for the implementation of a circular flow bioeconomy. Consequently, huge promise lies in their tailored optimization in order to produce fine chemicals, small molecules and biopharmaceuticals with high yield and quality. This optimization includes the modification of genes through beneficial mutations that modulate enzyme activity or full knockouts of product-degrading enzymes (proteases, lipases, nucleases), as well as metabolic engineering of complete heterologous biosynthetic pathways (Li et al., 2015).

Historically, microbial expression hosts were optimized by random mutagenesis and screening for mutant strains with favorable production characteristics. Later, homology-directed recombineering techniques enabled targeted, more precise genome modifications (Datsenko and Wanner, 2000; Orr-Weaver et al., 1981). However, these systems require antibiotic or auxotrophic marker selection, leave recombinase scar sequences after marker excision, and involve considerable labor.

Recently, the use of RNA-guided nucleases has revolutionized genome engineering and enabled the highly efficient and precise modification of genomes (Jinek et al., 2012; Zetsche et al., 2015) almost at will in a broad variety of organisms including bacteria (Jiang et al., 2013), microbial eukaryotes such as yeast (DiCarlo et al., 2013) up to mammals (Shen et al., 2013).

Multiple Cas9-based systems for genome modification of *Escherichia coli* drastically facilitated genome editing and enabled almost arbitrary genome modifications even without antibiotic marker genes and recombinase scars (Jiang et al., 2013; Reisch and Prather, 2015). These tools are thus in principle ideal for optimizing strains for industrial applications but are considered to be part of an increasingly complex intellectual property landscape (Ledford, 2022). An alternative to Cas9 are the members of the Cas12a family, which combine the DNA endonuclease activity with advantages of additional RNase activity to process their own guide RNA (gRNA) and a smaller protein size compared to Cas9.

Here, we describe an efficient genome editing system for *E. coli* based on the Cas12a-like nuclease MAD7 in combination with a λ-Red recombination system (Datsenko and Wanner, 2000). We characterize the system with different donor DNA formats (ssDNA and dsDNA) and realize different modifications of multiple gene loci. Finally, we combine the modular design of the system with state-of-the-art donor DNA assembly to introduce two glucose-uptake altering modifications, thereby creating an *E. coli* strain with improved plasmid production. Altogether, our approach provides a flexible genome editing system that enables fast and efficient genetic engineering to optimize *E. coli* strains.

## Results

### A genome editing toolkit based on MAD7 nuclease

MAD7 (also referred to as *ErCas12a*) is a Cas12a-like nuclease from *Eubacterium rectale* that cleaves its target DNA efficiently after 5’-YTTN-3’ protospacer adjacent motives (PAM, Figure 1 A). Because MAD7 “is royalty-free for both academic and commercial research and development use” (Inscripta, 2021; Price et al., 2020; Rojek et al., 2022) it serves as ideal tool for strain engineering in an industrial and commercial environment. MAD7-mediated genome engineering has been demonstrated both in eukaryotic systems including fungi (Jarczynska et al., 2021), plants (Lin et al., 2021), vertebrate and mammalian systems (Liu et al., 2020; Wierson et al., 2019), as well as in the prokaryotic organisms *E. coli* and *Bacillus subtilis* (Price et al., 2020). Previous use of MAD7 in *E. coli* focused on screening or nuclease engineering (Dewachter et al., 2022; Liu et al., 2019). Motivated by these reports, we decided to design an efficient CRISPR-MAD7 toolkit optimized for the application of genetic engineering in *E. coli* in an industrial setting.

**Figure 1:**
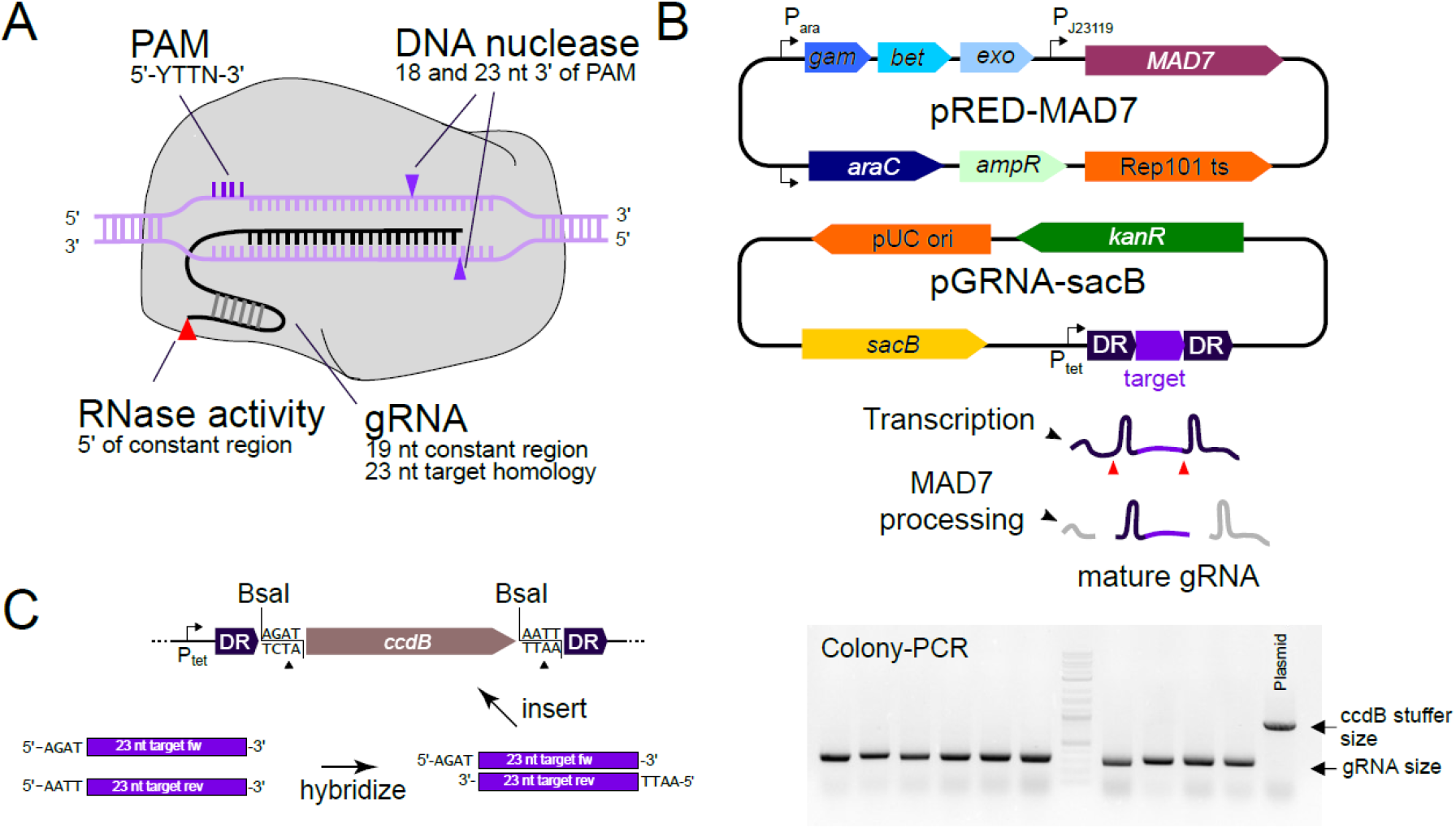
Design of the CRISPR-MAD7 system. **A)** Activity of MAD7 nuclease. MAD7 recognizes guide RNAs containing a constant direct repeat region, a 5’-YTTN-3’ protospacer adjacent motif (PAM) and a 23 nt variable target region, and cleaves the DNA 18 and 23 nucleotides (nt) 3’ of the PAM. In addition, MAD7 processes the RNA by cleaving 5’ upstream of the direct repeat. **B)** Design of CRISPR-MAD7 plasmids. The pRED-MAD7 activity plasmid combines the arabinose-inducible λ Red system (*gam*, *bet*, *exo*) with constitutively expressed MAD7. The pGRNA-sacB specificity plasmid contains the constitutively expressed pGRNA-transcript. **C)** pGRNA cloning. The vector pGRNA-sacB-ccdB is digested using BsaI. Then, oligonucleotides consisting of 5’ AGAT followed by the forward target sequence are hybridized to oligonucleotides with 5’ AATT followed by the reverse complement of the target sequence, yielding a dsDNA fragment with compatible overhangs for ligation into pGRNA-sacB with extremely high efficiency, as shown by a representative colony PCR for pGRNA-sacB-fruR.

### A modular plasmid system to separate activity and specificity

In *E. coli*, homology-directed gene targeting using linear DNA templates relies on activity of the λ Red system (Datsenko and Wanner, 2000), which therefore needs to be co-expressed alongside MAD7 and the guide RNA. Here, we designed the CRISPR-MAD7 system as a set of two expression plasmids. All activity-related components (MAD7 and λ Red system) are expressed from a single plasmid (pRED-MAD7, Figure 1 B), that carries a temperature-sensitive origin of replication (ori). For this, we inserted a MAD7 expression cassette to the λ Red plasmid pKD46 (Datsenko and Wanner, 2000). There, MAD7 expression is driven by the constitutive synthetic J23119 promoter (Anderson, iGEM parts), while the λ Red system is expressed from an inducible araBAD promoter, offering temporal control over recombination activity inside the cell. Furthermore, the temperature-sensitive ori enables curing the plasmid at elevated temperatures.

The specificity-mediating gRNA(s) are expressed from a second plasmid, pGRNA-sacB, which is based on a pUC backbone. A tet-promoter drives constitutive transcription of a MAD7 guide RNA cassette consisting of a MAD7-specific direct repeat sequence (DR) followed by the target sequence/protospacer and a second DR. Both DR sequences are processed by MAD7, releasing a gRNA with defined 5’ and 3’ ends (Figure 1 B, bottom). In addition to the gRNA, pGRNA-sacB contains the levansucrase-encoding *sacB* gene from *B. subtilis* to enable plasmid curing (Gay et al., 1985) in sucrose medium.

To enable rapid cloning of pGRNA plasmids for any desired target sequence, we designed the vector pGRNA-sacB-ccdB containing the ORF of the CcdB toxin (Bernard et al., 1994) flanked by recognition sequences for the type IIS restriction enzyme BsaI. This enables insertion of a target sequence by Golden-Gate cloning (Engler et al., 2008), thereby replacing the toxic *ccdB* sequence and ensuring that only colonies containing the correctly assembled pGRNA plasmid are obtained with high efficiency (Figure 1 C). The 27-mer oligonucleotides required for this cloning approach are commercially available from a wide range of vendors with short delivery times, which meets the industrial needs to be fast and to rely on multi-sourcing.

Taken together, our MAD7 system is designed for robust and rapid genome engineering in *E. coli*. Due to its high efficiency, our pGRNA cloning pipeline is compatible with an automated high throughput cloning strategy to generate large numbers of pGRNA plasmids with minimal hands-on time.

### MAD7 is active and allows genome-editing in E. coli

Next, we tested the activity of our MAD7 system in *E. coli*. It has been previously reported that repetitive DNA cleavage by constitutive Cas-nuclease activity induces cell death in *E. coli*, which can be used to select for cells engineered in the sequence target region or PAM (Figure 2 A) (Jiang et al., 2013; Reisch and Prather, 2017, 2015). Therefore, we first tested whether MAD7-gRNA complex formation and activity could be detected when pRED-MAD7 is expressed alongside a pGRNA plasmid carrying a previously reported gRNA sequence targeting the *galK* locus (pGRNA-galK) (Liu et al., 2019). To this end, an *E. coli* K12-derived strain bearing pRED-MAD7 was transformed with equimolar amounts of pGRNA-galK or a control plasmid. We detected a 10^4^-10^5^-fold reduction of colony-forming units (cfu) with the *galK* gRNA compared to the construct without gRNA expression cassette (Figure 2 B). Since cells are killed by persistent nuclease activity, this indicates that MAD7 is indeed active when supplied with a proper gRNA, thereby confirming the suitability of our plasmid design.

**Figure 2:**
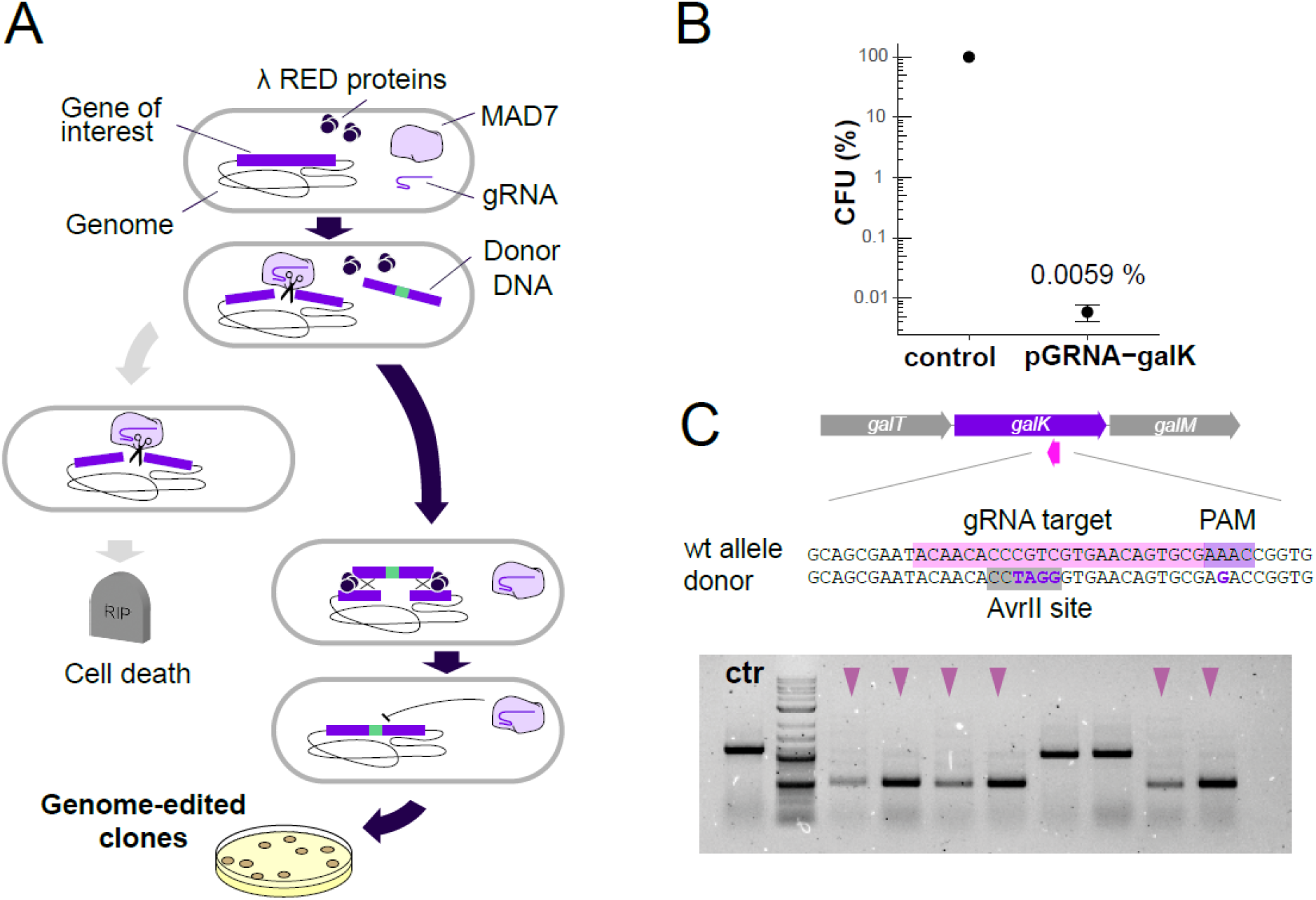
Principle of MAD7-mediated genome editing. **A)** Necessary components for genome engineering. When MAD7 and a guide RNA are present, MAD7 induces double strand breaks at the gene of interest, inducing cell death. When a donor DNA is additionally present, the λ Red system mediates homology-directed repair, introducing the desired genome edit (green segment). Only the edited gene of interest is no longer recognized by MAD7, so that only genome-edited clones survive. **B)** MAD7 activity. Cells were transformed with both pRED-MAD7 as well as equimolar amounts of either pGRNA-galK containing a gRNA targeting the gene *galK*, or a control plasmid without a gRNA. Surviving cells (colony forming units, cfu) were quantified after plating on LB-agar with kanamycin and ampicillin, revealing a decrease to less than 0.01% when MAD7 cleaves *galK*. **C)** MAD7-mediated genome editing of galK. The same gRNA was used as in B), but the pGRNA plasmid additionally contained a donor DNA sequence that introduced an AvrII restriction site. Representative gel shows identification of positive clones by colony PCR and AvrII digest. 8 of 15 tested clones shown.

After demonstrating the system’s activity, we next tested whether we were able to combine it with the λ Red activity to manipulate the genome. We designed a donor DNA to mutate the gRNA recognition site and thereby introduce a restriction site for AvrII into the *galK* gene (Figure 2 C), which enables restriction enzyme-based genotyping for identification of engineered clones. We cloned the donor DNA into pGRNA-galK and transformed this construct into pRED-MAD7 containing cells in which λ Red expression had been induced. We analyzed single clones by colony PCR followed by AvrII digest and found that most (12 of 15) clones contained the AvrII site, indicating correct genome editing (Figure 2 C).

### Simple and efficient genome editing using ssDNA oligonucleotides as donor DNA

While including the donor DNA sequence on the pGRNA plasmid was suitable for genome editing, it requires substantial hands-on time for cloning of each individual genome edit. Thus, we next tested if we could omit this cloning step to save time and resources, as well as increase flexibility, by directly suppling donor as commercially available single-stranded DNA (ssDNA) oligonucleotides. We designed an ssDNA donor consisting of 50 bp homologies spaced 300 bp apart in the *galK* gene, ca. 150 bp upstream and downstream of the gRNA target site. We co-transformed pGRNA-galK and varying amounts (500 pmol – 5 nmol) of ssDNA oligo donor and measured the fraction of genome edited clones by colony PCR. This experiment revealed that when sufficient ssDNA oligo is present, this simplified genome editing approach is highly effective, yielding up to 100 % positive clones when 5 nmol donor DNA were used (Figure 3 A).

**Figure 3:**
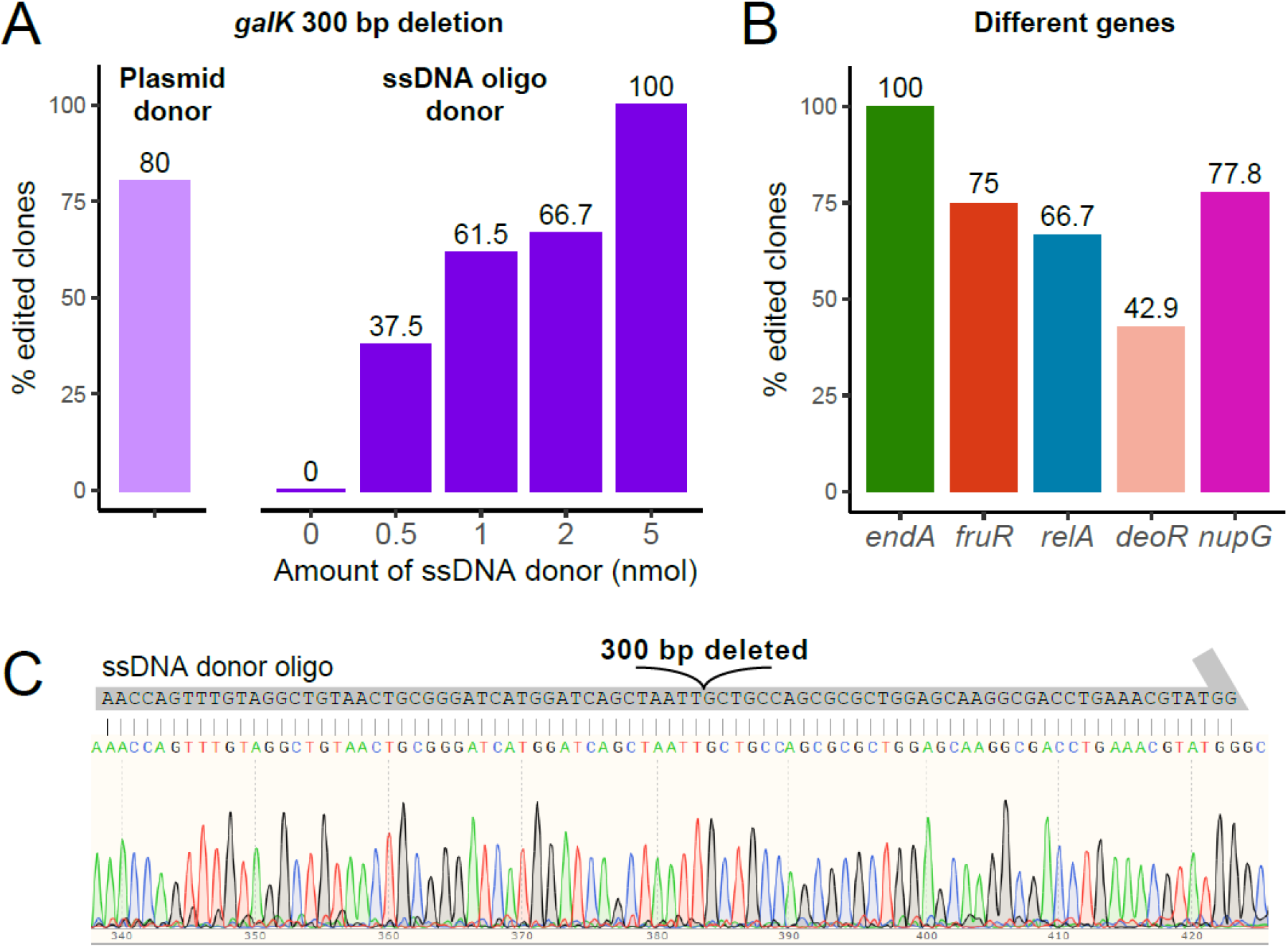
Efficient genome editing using ssDNA oligos as donors. **A)** Titration of donor oligo. Cells were transformed with pGRNA-galK alongside the indicated amounts of ssDNA oligo to delete 300 bp within the *galK* gene. Efficiency was quantified by colony PCR. Efficiency of *galK* editing (Figure 2) shown for comparison. **B)** Efficiency of deleting different genes. pGRNA plasmids for 5 different genes were transformed alongside 5 nmol of the respective ssDNA oligos, and editing efficiency was quantified using colony PCR. **C)** Representative Sanger sequencing of edited clone. The *galK* locus from a successful clone in (A) was amplified using colony PCR and analyzed by Sanger sequencing, highlighting that the deletion occurred with 100 % accuracy.

We showed that using our system, the *galK* gene can be modified accurately and efficiently. Thus, we next tested the general efficiency of MAD7-mediated genome editing by deleting five genes in different loci of the *E. coli* genome. We therefore cloned pGRNA plasmids targeting *endA*, *fruR*, *relA*, *deoR* and *nupG*, and designed ssDNA deletion donors (650 – 2243 bp deletion). We achieved efficient editing with more than 50 % positive clones for all targets except for *deoR* (Figure 3 B), when cells were recovered for 2 h (Supplementary Figure 1). Moreover, Sanger sequencing of the colony PCR products of *galK* revealed that genome editing was 100 % accurate in positive clones (Figure 3 C).

Taken together, we have demonstrated that our toolkit for MAD7-based genome editing is ideally suited for flexible, time- and resource efficient genome engineering in *E. coli* with high accuracy.

### Engineering complex genotypes by sequential editing and plasmid curing

Strain engineering often requires modifications or deletions in multiple genomic loci to achieve the genotype that confers the desired strain properties. In an industrial context, this for instance includes metabolic engineering to maximize yield or deleting multiple genes to avoid unwanted product degradation or impurities.

Our MAD7 system is ideally suited for this purpose, as the two plasmids pRED-MAD7 and pGRNA can be cured independently from each other. This allows for in principle unlimited numbers of sequential edits by iterating pGRNA/DNA donor transformations, selection of positive clones and pGRNA curing, while maintaining pRED-MAD7 (Figure 4 A). Once all desired genome modifications are obtained, pRED-MAD7 is cured from the strain, thus leading to a plasmid-free base strain. This step is crucial for industrial applications to enable transformation of product-encoding plasmids.

**Figure 4:**
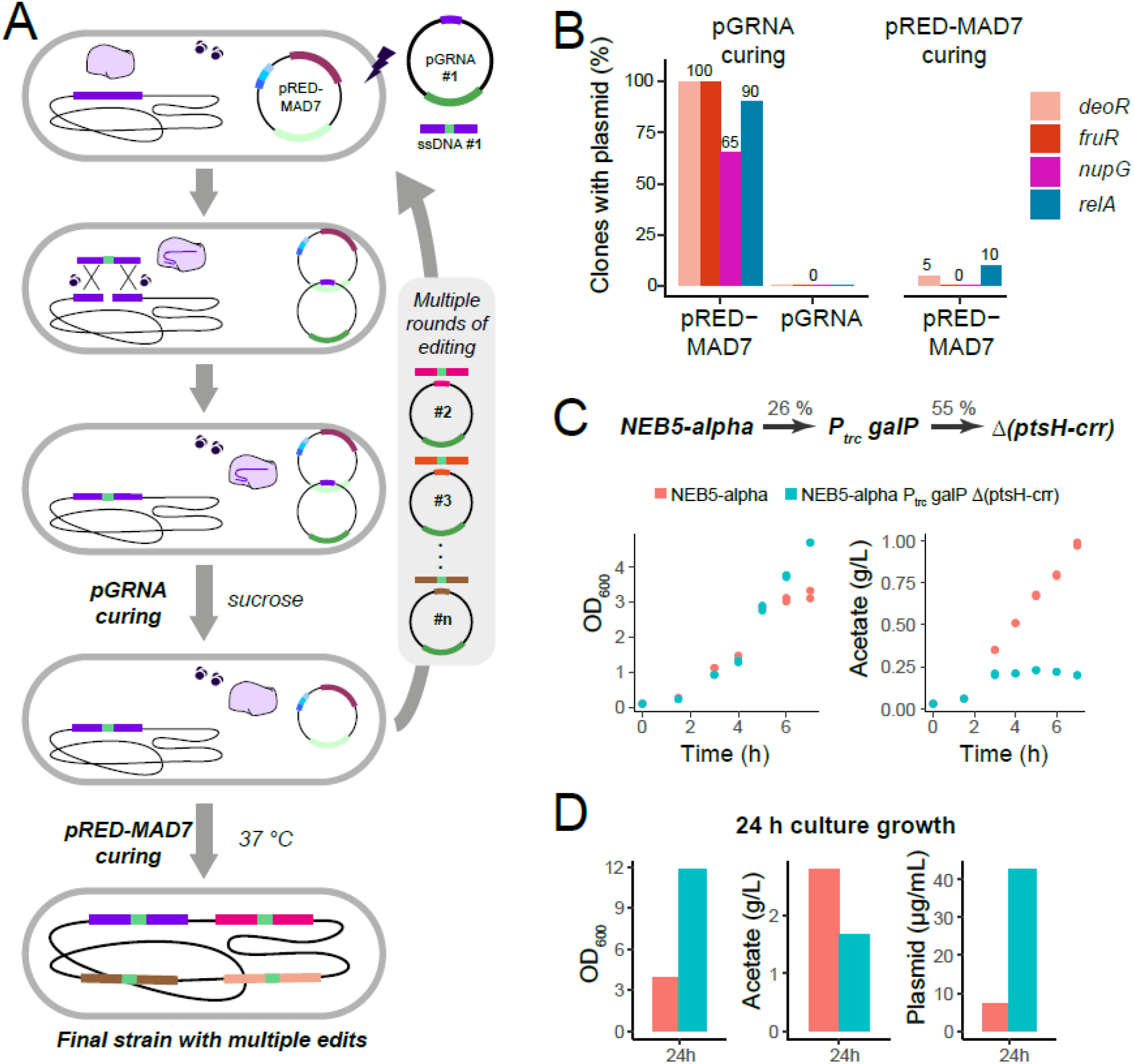
Sequential editing of multiple loci using MAD7-CRISPR. **A)** Workflow of multiple edits. Cells are transformed with pRED-MAD7, and λ Red expression is induced. Then pGRNA and donor DNA is transformed to achieve genome editing. In a correct clone, pGRNA is then cured by plating on agar plates containing sucrose. At this point, an arbitrary number of additional genome editing rounds can be performed. Finally, pRED-MAD7 is cured at 37 °C, yielding the plasmid-free strain with the desired genome edits. **B)** Plasmid curing is efficient and orthogonal. For 4 different edited strains, pGRNA plasmids were cured first, followed by curing of pRED-MAD7. **C)** Creation of an NEB5-alpha strain with reduced overflow metabolism NEB5-alpha cells were edited in two steps to yield NEB5-alpha *P*_*trc*_*-galP Δ(ptsH-crr)*. The efficiency of each editing step is indicated. The novel strain exhibits strongly reduced acetate formation, indicating reduced overflow metabolism. **D)** Novel NEB5-alpha strain shows increased plasmid productivity. NEB5-alpha *P*_*trc*_*-galP Δ(ptsH-crr)* harboring a model plasmid (pUC ori, kanamycin resistance) was cultured overnight, and showed a dramatic plasmid yield increase per culture volume.

We tested the efficiency of curing each plasmid individually and found that the first step of eliminating pGRNA was extremely efficient, as evident from a complete loss of kanamycin resistance for four different pGRNA plasmids (Figure 4 B). We even noticed that up to 30 % of clones had spontaneously lost pRED-MAD7 already during pGRNA-curing, potentially saving the second curing step. Afterwards, also pRED-MAD7 could be cured with very high efficiencies of 90-100 %, indicating that sequential genome editing can be achieved reliably with our MAD7 system.

We decided to demonstrate sequential editing by generation of an *E. coli* strain with an alternative glucose uptake system. In *E. coli*, acetate accumulation by overflow metabolism can be limiting (Anda et al., 2006). Deletion of the phosphotransferase system (PTS) genes *ptsH*, *ptsI* and *crr* alongside constitutive expression of the galactose permease-encoding *galP* gene circumvents this problem by ATP-based glucose import (Knabben et al., 2010). As a specific showcase, we engineered a metabolically improved strain based on the commonly used NEB5-alpha by introducing these two modifications.

### Resource-optimized genome editing for industrial applications

To improve the throughput and resource efficiency of our system even further, we decided to use benchtop DNA assembly to produce the donor DNA for our MAD7 system directly in the lab. We designed a trc-promotor (P_trc_) (Brosius et al., 1985) region replacing the endogenous promoter of *galP* flanked by 500 bp homology arms for the first genome editing step, and donor consisting of 500 bp homologies 5’ of *ptsH* and 3’ of *crr* for the second step.

We then synthesized these DNA fragments on a BioXp™ System (Codex DNA, Inc., San Diego, CA), and directly transformed the output DNA material alongside the pGRNA plasmids into NEB5-alpha cells without any further purification. The first step of replacing the *galP*-promoter was successful in 26 % of the analyzed clones (Figure 4 C). After curing a successfully modified *P*_*trc*_*-galP* clone from pGRNA-sacB-PgalP, we again induced the λ Red system and co-transformed pGRNA-sacB-ptsH with the second BioXp™-assembled donor, yielding the desired PTS deletion in 55 % of the analyzed clones.

Next, we tested how the new NEB5-alpha *P*_*trc*_*-galP Δ(ptsH-crr)* strain performed under relevant conditions by measuring acetate accumulation over time during shake flask batch cultivation (LB supplemented with 10 g/L glucose). While the unmodified NEB5-alpha strain continuously accumulated acetate up to 1 g/L within 6 h of cultivation, acetate in NEB5-alpha *P*_*trc*_*-galP Δ(ptsH-crr)* did not exceed 0.25 g/L within the first 6 h, despite a comparable or even increased cell density (Figure 4 C). When grown to saturation, NEB5-alpha *P*_*trc*_*-galP Δ(ptsH-crr)* had an almost four-fold increased cell density when compared to NEB5-alpha, at about half the acetate production (Figure 4 D). This clearly indicated that the combined genome edits successfully changed the metabolic profile of NEB5-alpha and re-routed the metabolic use of the available carbon source away from overflow metabolites towards increased productivity of biomass.

Motivated by these results, we next tested whether these modifications also improved the plasmid production, which is a common laboratory and industrial application of NEB5-alpha. We transformed NEB5-alpha *P*_*trc*_*-galP Δ(ptsH-crr)* with a model plasmid comprising a pUC ori and a kanamycin resistance gene. Indeed, when cultured under identical conditions as before, plasmid yield per mL culture volume was increased by a factor of 6 (Figure 4 D). This demonstrates the versatility and efficiency of our CRISPR-MAD7 system and highlights the potential to combine several genomic edits to generally improve *E. coli* production strains, as well as tailor them towards specific needs of industrial applications.

## Discussion

### CRISPR-MAD7 allows robust genome editing

CRISPR-based genome editing is widely used in academic research, but the intellectual property situation has been described as complex (Ledford, 2022). This has potentially limited its adoption in the biotechnological and pharmaceutical industry. Here, we present a MAD7 nuclease-based system for efficient and rapid genome engineering of *E. coli* cells with a special focus on reduced hands-on time and increased throughput, which are general requirements in an industrial research and development environment.

We demonstrate that our CRISPR-MAD7 system allows robust gene deletion and editing in *E. coli* (Figures 2-4). Using oligo or plasmid-coded donor DNA, we achieve efficiencies ranging from 43-100 %, approaching efficiencies reported for Cas9-based (ca. 85 – 100 %, (Reisch and Prather, 2015)) and Cpf1-based genome editing (ca. 81 % - 100 %, (Yan et al., 2017)) in *E. coli*. Lower efficiencies potentially result from different gRNA affinities to the respective loci or varying MAD7 processing efficiency of the direct repeats in the gRNA cassette. Other explanations could be gene-specific editing propensities due to chromosome structure, locus position or methylation.

### Our CRISPR-MAD7 system is compatible with high-throughput genome editing

When developing our CRISPR-MAD7 system, we prioritized speed and ease-of-use. In combination with benchtop DNA assembly via BioXp™, genome editing is reduced to a one-stop procedure between *in silico* target design and *in vivo* editing. We are thereby able to dramatically shorten the design cycle between benchmarking a microbial strain in its desired application, identifying the need for further strain improvements, and realizing additional genetic edits. This establishes a new paradigm for strain engineering as powerful tool to address challenges and bottlenecks in industrial microbial production processes.

The throughput of CRISPR-MAD7 can be increased through several routes. MAD7 can process an array of gRNAs to simultaneously target multiple gene loci, thus allowing gRNA multiplexing (Lin et al., 2021). In addition, we designed our CRISPR-MAD7 system with automation in mind. Due to *ccdB*-mediated counter-selection, pGRNA plasmids can be cloned automatically in 96-well plates. With additional automation tools such as high-throughput electroporation and automated colony picking and analysis, we envision that the throughput of CRISPR-MAD7 can be increased by orders of magnitude.

Taken together, we are convinced that our CRISPR-MAD7 toolkit represents a major advance that allows industrial research and development to harness the power of CRISPR genome engineering, and to create novel microbial cell factories for next-generation biologics in a robust, time- and cost-efficient manner.

## Materials and Methods

### Plasmid construction

A DNA fragment encoding the MAD7 ORF including regulatory sequences (J23119 promoter and T7-terminator) and 5’ and 3’ overlaps with adjacent sequences of the NcoI site in pKD46 was synthetized by GeneArt (Thermo Fisher Scientific). pRED-MAD7 was constructed by integration of this fragment into NcoI-linearized pKD46 (Datsenko and Wanner, 2000) by Gibson assembly (New England Biolabs).

To construct pGRNA-galK, the gRNA transcription cassette including tet-Promoter, spacer sequence, 21 nt MAD7-specific direct repeat, 23 nt *galK* target sequence protospacer, 19 nt MAD7-specific direct repeat and *rnpB* terminator, was synthetized as two genestrings by GeneArt (Thermo Fisher Scientific) and assembled into PCR-linearized (primers 454 and 455) pUC19-Kan by Gibson assembly (New England Biolabs).

pGRNA-galK-donor was created by assembly of PCR-linearized pGRNA-galK (primers 488 and 489), with oligo 477 by Gibson assembly (New England Biolabs).

pGRNA-sacB-galK was constructed by introduction of *sacB* into pGRNA-galK. A DNA fragment encoding the SacB ORF including the *P*_*cat*_ promoter and the ribosome binding site from bacteriophage T7 gene 10 (Olins & Rangwala, 1989) and was created by gene synthesis (GeneArt, Thermo Fisher Scientific). pGRNA-galK was linearized by PCR using primers 95 and 391 and assembled with the *sacB*-fragment by Gibson assembly (New England Biolabs).

For construction of pGRNA-sacB-ccdB, pGRNA-sacB-galK was PCR-amplified using primers 491 and 492 and the resulting PCR-product was assembled with oligo 550, oligo 551 and a gene string encoding the *ccdB* stuffer (GeneArt) using Gibson assembly (New England Biolabs).

pGRNA-sacB-endA was cloned by PCR-amplification of pGRNA-sacB-ccdB using primers 542 and 543 and subsequent circularization of the PCR product by Gibson assembly (New England Biolabs).

All other pGRNA constructs were cloned as described in Fig. 1 C. Respective forward and reverse oligonucleotides (10 pmol each) were hybridized in 50 μL annealing buffer (50 mM NaCl, 10 mM Tris, 1 mM EDTA) by heating to 95 °C for 5 min followed by 0.1 °C/s cooling to 4 °C in a thermocycler. 1 μL of ds-oligo solution was ligated into 50 ng BsaI-digested pGRNA-sacB-ccdB using T4 DNA Quick Ligase (New England Biolabs).

Sequences of the oligonucleotides are listed in Supplemental Table 1. All plasmids are listed in Supplemental Table 2. The full sequences of all plasmids are included in the supplemental information.

### Target and donor design and synthesis

Target sequences were designed based on the genome sequence of *E. coli* MG1655 (Blattner et al., 1997). Target sequences were identified in close proximity to the desired edit and 3’ of a 5’-YTTN-3’ PAM, with GC content close to 50 %. Donor DNA was designed such that either the PAM or the target region was altered or deleted after the edit. Single stranded donor DNA was synthetized as oligonucleotides by Thermo Fisher Scientific (Waltham, MA, USA). Double stranded donor DNA was assembled from pooled oligos using the BioXp™ 3250 system (Codex DNA, Inc., San Diego, CA).

### Genome modification

Electrocompetent cells of the strain to be edited were prepared as previously described (Woodall, 2003), transformed with pRED-MAD7 by electroporation (2 mm cuvette, 2.5 kV, 25 μF, 200 Ω) and a single clone was selected from LB-Agar containing 100 μg/mL Ampicillin. The clone was cultured overnight in 5 mL SOB medium (0.5 % yeast extract, 2 % Phytone, 10 mM NaCl, 2.5 mM KCl, 10 mM MgCl_2_, 10 mM MgSO_4_) containing 100 μg/mL Ampicillin (SOB-Amp) at 30 °C and 220 rpm in a 14 mL round bottom tube. 100 mL SOB-Amp were inoculated to an OD_600_ of 0.05 with overnight culture and incubated at 30 °C and 220 rpm. When the culture reached an OD_600_ of 0.3 – 0.4, λ Red expression was induced by addition of 6.5 mL 1 M arabinose solution for 45 min. The culture was cooled for 10 min on ice. Cells were pelleted by centrifugation at 3000 x g rcf and 4 °C for 10 min and washed three times with sterile 4 °C cold H_2_O. Cells were washed in 2 mL cold 10 % glycerol and resuspended in 0.8 mL 10 % glycerol, of which 50 μL aliquots were stored at −80 °C. 100 – 250 ng pGRNA and donor DNA were co-transformed by electroporation (2 mm cuvette, 2.5 kV, 25 μF, 200 Ω) and cells were recovered in 1 mL SOC medium (New England Biolabs, #B9020) at 30 °C and for 90 min. Cells were pelleted again, resuspended in 2 mL LB containing 100 μg/mL Ampicillin and 50 μg/mL Kanamycin and incubated at 30 °C 220 rpm in a 14 mL round bottom tube for 2 h. 20 μL and 200 μL of the culture were plated on LB-Agar containing 100 μg/mL Ampicillin and 50 μg/mL Kanamycin and plates were incubated for > 24 h at 30 °C. We noticed a decrease in editing efficiency if cells were incubated overnight instead of 2 h before plating (Supplemental Figure 1), hinting at mutants arising over time that evade Mad7-mediated DNA cleavage.

All strains created in this study are listed in Supplemental Table 3.

### Transformant counting

Agar plates were scanned using a flatbed scanner (Epson V750 Pro) and colonies in digital images were counted using Fiji/ImageJ’s Multi-point tool (Schindelin et al., 2012). Data was analyzed and visualized using R (R Core Team, 2022) and ggplot2 (Wickham, 2016).

### Quantification of editing

For quantification of modification editing (insertion of AvrII site in *galK*), single clones were analyzed by colony PCR (DreamTaq Mastermix, Thermo Fisher Scientific) using primers 478 and 479. 0.5 μL AvrII (New England Biolabs) was added to each reaction, mixed by pipetting up and down, incubated for 1 h at 37 °C. DNA fragments were analyzed by agarose gel electrophoresis. Correct edits were identified by appearance of a 0.5 kb band and validated by Sanger sequencing. Deletion edits were quantified by colony PCR (DreamTaq Mastermix, Thermo Fisher Scientific) using the primers for the edited locus as listed in Supplemental Table 1. PCR products were analyzed by agarose gel electrophoresis and success was judged by size of the product band.

### Plasmid Curing

To cure pGRNA, cells were singled out on LB-Agar containing 100 g/L sucrose and incubated at 30 °C for >24 h. Single colonies were sequentially replicated on LB-agar plates containing either Ampicillin (100 μg/mL), Kanamycin (50 μg/mL) or no antibiotic (as control). Plates were incubated overnight at 30 °C and plasmid presence was scored by growth on the respective antibiotic. To cure pRED-MAD7, cells were singled out on LB-Agar and grown overnight at 37 °C. Single colonies were sequentially replicated on LB-agar plates containing either Ampicillin (100 μg/mL) or no antibiotic. Plates were incubated overnight at 37 °C and presence of pRED-MAD7 was scored by growth on LB-Ampicillin.

### Quantification of overflow metabolism and plasmid productivity

Cells were cultured in 100 mL LB with additional 10 g/L glucose and 50 μg/mL kanamycin in a 300 mL Erlenmeyer flask. Flasks were inoculated from overnight cultures to an OD of 0.1. The cultures were then sampled every 1-2h by measuring OD as well as collecting 2 mL of culture in an Eppendorf tube, followed by centrifugation, and transferring 800 μL of cell-free supernatant to a fresh sample tube. Acetate concentration in the supernatant was measured using the Cedex Acetate V2 Bio HT kit on a Cedex HT system (Roche Custom Biotech). OD_600_ was measured using a Ultrospec 2100 Spectrophotometer (Amersham Biosciences).

For plasmid concentration measurements, cells were inoculated from an overnight culture to an OD_600_ of 0.1 in 10 mL LB with appropriate antibiotics and additional 10 g/L glucose in a 100 mL Erlenmeyer flask at 37 °C and 220 rpm for overnight. A cell mass of an equivalent of a OD_600_ 3 was used for a DNA mini preparation using a QIAprep® Spin Miniprep Kit (Qiagen). DNA was eluted in 50 μL elution buffer and measured using a NanoDrop one (Thermo Fisher Scientific).

## Acknowledgments

We thank Natalie Schmidt for support with Acetate measurements. We thank Andreas Matern, Fabian Bindel, Elena Antonov, Sebastian Scholz, Angel Corcoles Garcia and Julius Kühn for critical reading and helpful comments on the manuscript. We thank the MP USP Molecular Biology Lab members for their support.

## Author contributions

M.M., W.W., C.S. performed experiments and analyzed data. D.D. and C.S. conceived the study. M.M., D.D. and C.S. supervised the study. M.M. and C.S. wrote the manuscript with input from the other authors.

