## Supplemental Information for "A MAD7-based genome editing system for *E. coli*"

#### Supplementary Figure 1

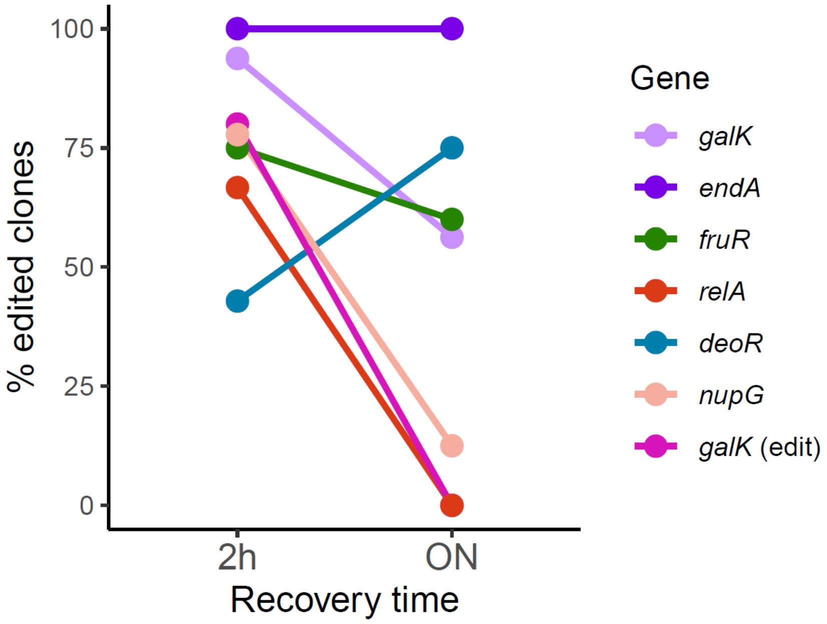


**Supplementary Figure 1: Genome editing efficiency is dependent on recovery time**. pGRNA plasmids for 5 different genes were transformed alongside 5 nmol of the respective ssDNA oligos to delete the indicated genes. Following recovery, cells were incubated in liquid medium either 2h or overnight before plating (compare material & methods). Editing efficiency was quantified using colony PCR, revealing that for most genes, editing efficiency substantially decreased with longer incubation times.

#### Supplemental Table 1: List of oligonucleotides used in this study

| ID | Name | Sequence (5‘ -> 3‘) | Comment |
|---|---|---|---|
| 95 | Backbone-1 | CCAGCAAAAGGCCAGGAAC | pGRNA-sacB-galK cloning |
| 284 | ilvH-R85-fw | GCAGGGCGCGCATGTTGAGCG | Colony PCR *fruR* |
| 285 | fruR-term-rev | GCTTGATGCCAGCGGGTTTGCGC | Colony PCR *fruR* |
| 301 | rsmE-A137-rev | GACCACACTGCTCACAGGCAGC | Colony PCR *endA* |
| 302 | yggI-fw | ATGAAAACATCCCGTCTCCCTATC | Colony PCR *endA* |

| | | | |
|---|---|---|---|
| 391 | Backbone-2-fw | TGCCATCGTATTAGCACCCCGCAC | pGRNA-sacB-galK cloning, Colony PCR fig. 1C |
| 421 | Backbone-3-rev | GACGCCGACGCTAACATCGAGCTG | Colony PCR fig. 1C |
| 454 | pUC-fw | CAGCTCGATGTTAGCGTCGGCGTCCCGGGAGCTGCATGTGTCAGAGG | pGRNA-galK cloning |
| 455 | pUC-rev | GTGCGGGGTGCTAATACGATGGCACTCGCTGCGCTCGGTCGTTCG | pGRNA-galK cloning |
| 477 | galK AvrII insertion Donor | CAGTAACTTCAAACGTACCCTGGTTGGCAGCGAATACAACACCTAGGG TGAACAGTGCGAGACCGGTGCGCGTTTCTTCCAGCAGCCAGCCCTGCGTG AT | Donor |
| 478 | galK-A47-fw | ACGGTTTCGTTCTGCCCTGCG | Colony PCR *galK* |
| 479 | galM-T29-rev | TCGCACCCCAGTCCATCAGCG | Colony PCR *galK* |
| 488 | galK-pUC fw | GCAGCCAGCCCTGCGTGATGTTGAGTGAGCTGATACCGCTCGC | pGRNA-galK cloning |
| 489 | galK-pUC-rev | CCAGGGTACGTTTGAAGTTACTGTTCAGAATCAGGGGATAACGCAGG | pGRNA-galK cloning |
| 491 | Backbone-2-rev | GTGCGGGGTGCTAATACGATGGCA | pGRNA-sacB-ccdB cloning |
| 492 | Backbone-3-fw | CAGCTCGATGTTAGCGTCGGCGTC | pGRNA-sacB-ccdB cloning |
| 505 | *galK* deletion 300 bp donor | AACCAGTTTGTAGGCTGTAACTGCGGGATCATGGATCAGCTAATTGCTGC CAGCGCGCTGGAGCAAGGCGACCTGAAACGTATGG | Donor |
| 519 | *relA* deletion Donor | TCGCGCGTTAAATAGTTGCGATTTGCCGATTTCGGCAGGTCTGGTCCCTA G GCCGAAATTTGCTCGTATCTACAATGTAGATTGATATATACTGTATCTA | Donor |
| 520 | mazF-Y5-rev | CAGATCGCCCATATCGGGTACG | Colony PCR *relA* |
| 521 | rlmD-L330-fw | AAGGCCAGCAGAATGCGCGTC | Colony PCR *relA* |
| 542 | endA-target-rev | CGGCTATCAGGTGCGCAAAAATGATCTACAAGAGTAGAAATTGGTAGC | Guide |
| 543 | endA-target-fw | CATTTTTGCGCACCTGATAGCCGAATTTCTACTCTTGTAGATAGCTGCC | Guide |
| 548 | *endA* deletion Donor | CTGGCTGATTGCATACCAAAACAGCTTTCGCTACGTTGCTGGCTCGTTTTT G AACGCATCGCGAAGGTGCAGGGCAATCATAACCCGTATGTGCAACGCG | Donor |
| 550 | ccdB-part-fw | CCCTTATACACAGCCAGTCTGCAGGTCTCGAATTTCTACTCTTGTAGATAG CT GCCACATCAAACTGGAGAATAAGGTCAGTTTCACCTGTTTTACGT | pGRNA-sacB-ccdB cloning |
| 551 | ccdB-part-rev | GACGCCGACGCTAACATCGAGCTGCTCGAGCCAAAAGTAAAAACCCGCC GA AGCGGGTTTTTACGTAAAACAGGTGAAACTGACCTTATTCTC | pGRNA-sacB-ccdB cloning |
| 552 | fruR-target-fw | AGATTGATCCCCGATCTGGAGAACACC | Guide |
| 553 | fruR-target-rev | AATTGGTGTTCTCCAGATCGGGGATCA | Guide |
| 557 | *fruR* deletion Donor | ATATTGATGTCCAGTCCCGTACTCTACGCGCCAGAGGGAAATTCACCTGG A CTGCTTTTATTCCAGGTTCCCACTGATTTCGCAAAAAAAGCCCAACGTG | Donor |
| 576 | relA-target-fw | AGATATTACCCAGGGGCGCGGTATTTC | Guide |
| 577 | relA-target-rev | AATTGAAATACCGCGCCCCTGGGTAAT | Guide |
| 592 | cysK-Y188-fw | CTGACTGGCGTCAGCCGCTAC | Colony PCR *ptsHI-crr* |
| 593 | pdxK-R146-fw | CCTGACCTTCCCGAAGCGTATCG | Colony PCR *ptsHI-crr* |
| 599 | P_galP target-fw | AGATCGTCGTACTCACCTATCTTAATT | Guide |
| 600 | P_galP target-rev | AATTAATTAAGATAGGTGAGTACGACG | Guide |
| 601 | metK-G290-fw | TGAATGGCTGACTTCTGCCACC | Colony PCR *galP* |

| | | | |
|---|---|---|---|
| 602 | galP-A37-rev | AATAAACGGCAGTGCGCCAGC | Colony PCR *galP* |
| 613 | ptsH-target-fw | AGATAAACTTTCGCCCCTCCTGGCATT | Guide |
| 614 | ptsH-target-rev | AATTAATGCCAGGAGGGGCGAAAGTTT | Guide |
| 642 | deoR-target-fw | AGATAAGAGTTGCCGGTAAAACACTGG | Guide |
| 643 | deoR-target-rev | AATTCCAGTGTTTTACCGGCAACTCTT | Guide |
| 644 | *deoR* deletion donor | CCGATCCGCAAAGGCTGGGTGCGTGACTGATCTCTGCTAAAAAGTGTAG TTGACGTATAACCGGATGACGTTTCGCGCCATCCGGTTATCAGAAGATTA A | Donor |
| 645 | ybjG-S124-fw | ATGCTGGCATCGCCTGTGGTCC | Colony PCR *deoR* |
| 646 | dacC-P344-fw | ACGGAACCGCAGCTTACCGCAC | Colony PCR *deoR* |
| 647 | speC-G630-fw | GCGCATAGCGCTTATATTCGCGG | Colony PCR *nupG* |
| 648 | mltC-G302-fw | CATCACCGCCTATAACGGCGGC | Colony PCR *nupG* |
| 649 | nupG-target-fw | AGATTAGCTCACTGGGTATCGCAGCGG | Guide |
| 650 | nupG-target-rev | AATTCCGCTGCGATACCCAGTGAGCTA | Guide |
| 653 | *nupG* deletion donor | GTTTGCAATTATTTGCCACAGGTAACAAAAAACCAGTCCGCGAAGTTGAT TATAAACACGTTCGTGTCCCGACAGGCACACAGACGGTTAGCCACTAATT | Donor |

### Supplemental Table 2: List of plasmids used in this study

| **Plasmid** | **Source** |
|---|---|
| pKD46 | (Datsenko and Wanner, 2000) |
| pUC19 | NEB/(Yanisch-Perron et al., 1985) |
| pUC-Kan | This study |
| pRED-MAD7 | This study |
| pGRNA-galK | This study |
| pGRNA-galK-donor | This study |
| pGRNA-sacB-galK | This study |
| pGRNA-sacB-ccdB | This study |
| pGRNA-sacB-endA | This study |
| pGRNA-sacB-fruR | This study |
| pGRNA-sacB-deoR | This study |
| pGRNA-sacB-relA | This study |
| pGRNA-sacB-nupG | This study |
| pGRNA-sacB-PgalP | This study |
| pGRNA-sacB-ptsH | This study |
| Plasmid with pUC backbone and $Kan^R$ marker | This study |

### Supplemental Table 3: List of strains

| **Strain** | **Source** |
|---|---|
| NEB5-alpha | New England Biolabs |
| K12 derivative | This Study |
| NEB5-alpha $P_{trc}$-galP | This Study |
| NEB5-alpha $P_{trc}$-galP ΔPTS (NEB5-alpha VH33) | This Study |

### Full Plasmid Sequences

**>pUC-Kan**

TTGAGATCCTTTTTTTCTGCGCGTAATCTGCTGCTTGCAAACAAAAAAACCACCGCTACCAGCGGTGGTTTGTTT
GCCGGATCAAGAGCTACCAACTCTTTTTCCGAAGGTAACTGGCTTCAGCAGAGCGCAGATACCAAATACTGTTC
TTCTAGTGTAGCCGTAGTTAGGCCACCACTTCAAGAACTCTGTAGCACCGCCTACATACCTCGCTCTGCTAATCC
TGTTACCAGTGGCTGCTGCCAGTGGCGATAAGTCGTGTCTTACCGGGTTGGACTCAAGACGATAGTTACCGGA
TAAGGCGCAGCGGTCGGGCTGAACGGGGGGTTCGTGCACACAGCCCAGCTTGGAGCGAACGACCTACACCG
AACTGAGATACCTACAGCGTGAGCTATGAGAAAGCGCCACGCTTCCCGAAGGGAGAAAGGCGGACAGGTATC
CGGTAAGCGGCAGGGTCGGAACAGGAGAGCGCACGAGGGAGCTTCCAGGGGGAAACGCCTGGTATCTTTAT
AGTCCTGTCGGGTTTCGCCACCTCTGACTTGAGCGTCGATTTTTGTGATGCTCGTCAGGGGGGCGGAGCCTAT
GGAAAAACGCCAGCAACGCGGCCTTTTTACGGTTCCTGGCCTTTTGCTGGCCTTTTGCTCACATGTTCTTTCCTG
CGTTATCCCCTGATTCTGTGGATAACCGTATTACCGCCTTTGAGTGAGCTGATACCGCTCGCCGCAGCCGAACG
ACCGAGCGCAGCGAGTCAGTGAGCGAGGAAGCGGAAGAGCGCCCAATACGCAAACCGCCTCTCCCCGCGCGT
TGGCCGATTCATTAATGCAGCTGGCACGACAGGTTTCCCGACTGGAAAGCGGGCAGTGAGCGCAACGCAATT
AATGTGAGTTAGCTCACTCATTAGGCACCCCAGGCTTTACACTTTATGCTTCCGGCTCGTATGTTGTGTGGAATT
GTGAGCGGATAACAATTTCACACAGGAAACAGCTATGACCATGATTACGCCAAGCTTGCATGCCTGCAGGTCG
ACTCTAGAGGATCCCCGGGTACCGAGCTCGAATTCACTGGCCGTCGTTTTACAACGTCGTGACTGGGAAAACC
CTGGCGTTACCCAACTTAATCGCCTTGCAGCACATCCCCCTTTCGCCAGCTGGCGTAATAGCGAAGAGGCCCGC
ACCGATCGCCCTTCCCAACAGTTGCGCAGCCTGAATGGCGAATGGCGCCTGATGCGGTATTTTCTCCTTACGCA
TCTGTGCGGTATTTCACACCGCATATGGTGCACTCTCAGTACAATCTGCTCTGATGCCGCATAGTTAAGCCAGC
CCCGACACCCGCCAACACCCGCTGACGCGCCCTGACGGGCTTGTCTGCTCCCGGCATCCGCTTACAGACAAGCT
GTGACCGTCTCCGGGAGCTGCATGTGTCAGAGGTTTTCACCGTCATCACCGAAACGCGCGAGACGAAAGGGC
CTCGTGATACGCCTATTTTTATAGGTTAATGTCATGATAATAATGGTTTCTTAGACGTCAGGTGGCACTTTTCGG
GAAGTTCCTATTCTCTAGAAAGTATAGGAACTTCGGAAATGTGCGCGGAACCCCTATTTGTTTATTTTTCTAAAT
ACATTCAAATATGTATCCGCTCATGAGACAATAACCCTGATAAATGCTTCAATAATATTGAAAAAGGAAGAGTA
TGATTGAACAAGATGGATTGCACGCAGGTTCTCCGGCCGCTTGGGTGGAGAGGCTATTCGGCTATGACTGGG
CACAACAGACAATCGGCTGCTCTGATGCCGCCGTGTTCCGGCTGTCAGCGCAGGGGCGCCTGGTTCTTTTTGTC
AAGACCGACCTCTCCGGTGCCCTGAATGAACTGCAAGACGAGGCAGCGCGGCTATCGTGGCTGGCCACGACG
GGCGTTCCTTGCGCAGCTGTGCTCGACGTTGTCACTGAAGCGGGAAGGGACTGGCTGCTATTGGGCGAAGTG
CCGGGGCAGGATCTCCTGTCATCTCACCTTGCTCCTGCCGAGAAAGTATCCATCATGGCTGATGCAATGCGGC
GGCTGCATACGCTTGATCCGGCTACCTGCCCATTCGACCACCAAGCGAAACATCGCATCGAGCGAGCACGTAC
TCGGATGGAAGCCGGTCTTGTCGATCAGGATGATCTGGACGAAGAGCATCAGGGGCTCGCGCCAGCCGAACT
GTTCGCCAGGCTCAAGGCGAGCATGCCCGACGGCGAGGATCTCGTCGTGACCCATGGCGATGCCTGCTTGCC
GAATATCATGGTGGAAAATGGCCGCTTTTCTGGATTCATCGACTGTGGCCGGCTGGGTGTGGCGGACCGCTAT
CAGGACATAGCGTTGGCTACCCGTGATATTGCTGAAGAGCTTGGCGGCGAATGGGCTGACCGCTTCCTCGTGC
TTTACGGTATCGCCGCTCCCGATTCGCAGCGCATCGCCTTCTATCGCCTTCTTGACGAGTTCTTCTGACTGTCAG
ACCAAGTTTACTCATATATACTTTAGATTGATTTAAAACTTCATTTTTAATTTAAAAGGATCTAGGTGAAGATCC
TTTTTGATAATCTCATGACCAAAATCCCTTAACGTGAGTTTTCGTTCCACTGAAGTTCCTATTCTCTAGAAAGTAT
AGGAACTTCGAGCGTCAGACCCCGTAGAAAAGATCAAAGGATCTTC

**>pRED-MAD7**

GTTGATGATACCGCTGCCTTACTGGGTGCATTAGCCAGTCTGAATGACCTGTCACGGGATAATCCGAAGTGGT
CAGACTGGAAAATCAGAGGGCAGGAACTGCTGAACAGCAAAAAGTCAGATAGCACCACATAGCAGACCCGCC
ATAAAACGCCCTGAGAAGCCCGTGACGGGCTTTTCTTGTATTATGGGTAGTTTCCTTGCATGAATCCATAAAAG
GCGCCTGTAGTGCCATTTACCCCCATTCACTGCCAGAGCCGTGAGCGCAGCGAACTGAATGTCACGAAAAAGA
CAGCGACTCAGGTGCCTGATGGTCGGAGACAAAAGGAATATTCAGCGATTTGCCCGATTGCGGCCGCAACCG
AGCTTGCGAGGGTGCTACTTAAGCCTTTAGGGTTTTAAGGTCTGTTTTGTAGAGGAGCAAACAGCGTTTGCGA
CATCCTTTTGTAATACTGCGGAACTGACTAAAGTAGTGAGTTATACACAGGGCTGGGATCTATTCTTTTTATCTT
TTTTTATTCTTTCTTTATTCTATAAATTATAACCACTTGAATATAAACAAAAAAAACACACAAAGGTCTAGCGGA
ATTTACAGAGGGTCTAGCAGAATTTACAAGTTTTCCAGCAAAGGTCTAGCAGAATTTACAGATACCCACAACTC
AAAGGAAAAGGACTAGTAATTATCATTGACTAGCCCATCTCAATTGGTATAGTGATTAAAATCACCTAGACCAA
TTGAGATGTATGTCTGAATTAGTTGTTTTCAAAGCAAATGAACTAGCGATTAGTCGCTATGACTTAACGGAGCA
TGAAACCAAGCTAATTTTATGCTGTGTGGCACTACTCAACCCCACGATTGAAAACCCTACAAGGAAAGAACGG
ACGGTATCGTTCACTTATAACCAATACGTTCAGATGATGAACATCAGTAGGGAAAATGCTTATGGTGTATTAGC
TAAAGCAACCAGAGAGCTGATGACGAGAACTGTGGAAATCAGGAATCCTTTGGTTAAAGGCTTTGAGATTTTC
CAGTGGACAAACTATGCCAAGTTCTCAAGCGAAAAATTAGAATTAGTTTTTAGTGAAGAGATATTGCCTTATCT
TTTCCAGTTAAAAAAATTCATAAAATATAATCTGGAACATGTTAAGTCTTTTGAAAACAAATACTCTATGAGGAT
TTATGAGTGGTTATTAAAAGAACTAACACAAAAGAAAACTCACAAGGCAAATATAGAGATTAGCCTTGATGAA
TTTAAGTTCATGTTAATGCTTGAAAATAACTACCATGAGTTTAAAAGGCTTAACCAATGGGTTTTGAAACCAAT
AAGTAAAGATTTAAACACTTACAGCAATATGAAATTGGTGGTTGATAAGCGAGGCCGCCCGACTGATACGTTG
ATTTTCCAAGTTGAACTAGATAGACAAATGGATCTCGTAACCGAACTTGAGAACAACCAGATAAAAATGAATG
GTGACAAAATACCAACAACCATTACATCAGATTCCTACCTACATAACGGACTAAGAAAAACACTACACGATGCT
TTAACTGCAAAAATTCAGCTCACCAGTTTTGAGGCAAAATTTTTGAGTGACATGCAAAGTAAGTATGATCTCAA
TGGTTCGTTCTCATGGCTCACGCAAAAACAACGAACCACACTAGAGAACATACTGGCTAAATACGGAAGGATC
TGAGGTTCTTATGGCTCTTGTATCTATCAGTGAAGCATCAAGACTAACAAACAAAAGTAGAACAACTGTTCACC
GTTACATATCAAAGGGAAAACTGTCCATACCCATGGCTTGTTCTCCAGCTGATGTGGCAGCTAAAAAAAACCCC
GCCCTGTCAGGGGCGGGGTTTTTTTTTCTCGAGTTAATTATTACAGATAGCGCTTGTTCTGGATGAAGTCGAAC
CAATCTTTGTTAGAAATTTTCAGTTTATCACGGGAGAATTTGCCATCCTCTTTCCAGTTTTCCGTGATTTGTTTGA
TCTCGTACAGGCCTTTCAGCGCGATACAGTACGCACCGTTAGCGTCCGCGTCTTTTGGCAGAGCATCACCGGCT
TTGGCGCTGTCATAAAAAATGTTGTTTTCGTTCAGAACCGGGCTGATCAGACGGTCGTAATCACGGTCTTCAAG
TTCAGACAGCGAGTTACGCATCTGTACGGTCAGACGAAAGATTTCGAAAATGTGCTGCACAATTTCGTAATCA
ATAATGTCCTGGCGCAGATCGTGGCCATCACGCCAGTTGATGTCAGTCATCTCCAGAGTCTTTTCCATATCTTTA
GTAATATCAATGGTATCGCTTTCGTTAGAAAAGCGACCGTTAACGAAACGGCGTTTGATGCGCACACCATAGG
TATACACAGACCAGCTAGATTTAGACATAACGGTGTTCTGAGTGATAAAGTTGTTGTAATCAAAGGTGAAACA
AAAGAGGTTCTTTTCAGAGTCATAGCGGATGCTGTCAAACTTCTTGATAAATTCACGTTTGGCATCCACGGTCA
GGTCTTTGAATTTGAAGATATTCACGAAACCAGTGGTCGGATCAATCTTGCTCGTGTAAGCTGCTGGAACGTAA
AAGATGCAGCCACACTGGTGGCCAACGTTTTTCAGTTTATCCGGGATATAAGTCAGTTGATAACCTTTCAGCAG
ACCGCCGTTCTCGGTGATGGAAATATCCTTAAATACCAGATAGTTTAATTTGTTGATCAGCATGGTTTCGAATTT
CTGGTACACTTGGCGTTCGACTTTGAAACGTCCTTTTTTGAAGCCGTAGCTCAAGTCTTCCATTGCGATGATAGC
GTTGTACTTGATAACCATTTTTGAAATTTCGTGGATCACCAGGCTCAAGTAACCTTCTTTGATCTCTTTGATCTTC
CCGATTTCCTTCCATTCTTTACGAGCGATCTGACGAGCACCTTCCTGCTGTTTCAGCTTGATCTGGTAGTCATAA
CCATTCACAATATTAAATGATTTCTGTTCAACGATGTTGCCGCAGGTGTCGATAACAGAAACGTAAATCAGGTT
GCGCTCGCCACGGTCAATACCAATCACATGCAGGTCTTTTTCTTTCGCGATATATTGAAGAATACGATCATTGAT
AAAGCCCGTTTTGTTAGCTTTAAAGTTAATGGTGATTGGCATGTGCAGGAAATATTTATCGTACGTATAACGGT
AGTCTTTTACAATGTTGGTAGCCGCTTCGTGGTGGCCAACGACATTTTTCAGTTTGGCGGCCTCATCAGACAGT
TCCTTATCAGATTTATCGTTGAAGTACTTATACAGCTCCTGGTAAATGTTTTCCGGAATGTTTTTGCGTACGATC
TGAATATTGCCAAACTGGTCTTTTTCCTCGGCTTCATAAGTGCGATTCACCAGGATAGAACCTTTTTTGTGGATG
ATCGGATTCTTGATGGAAGATTTGCGGAAGAAGATTTCCGCTTCGCCGTTCAGTTTCAGAACAATGTCTTTCAG

ATTTTCCTCGCTGAACAGGTTTTTCAGGTACATGGTGTGCAGGTTGTCATTGCCAGTCGATTTTTTGCTGAAGTC
TTTATTATAGATTTGAAACAGGTATAACTGACCCTTTTCCTGAAGCAGGTCAATGTCTTTTTCGCTGATATAGGT
CCAATCGATTTTGTAGCCCTGCAGTTCAACTTCGCGGTAGAAACCGCTGATGTCTTCGTACGTGGAAGTGTCGC
TGAAATCAAAGCCGAAATTTTTCCACTCCGGGTGAATCGCGATGCAGTTCTTGAAATAATCGATCAGGTCGTG
GCAGAAGGTGATGTCAAAGTCTTTGGAAGATTTGATATGTTTGTTCTGTTTATAGCCTTCCAGGATATACGCAG
ACGGTTTGTAGGTTTCTACACCTGTTTTGCTAGACAGGAATACTTTCGGGATCATTTTGTTCGGGCCCGGCAGC
AGGTTATAGATCATTTTTTTATAATCGCCTTTGTTTTCGGATGTGTTACCTTCAATGATTTTTTTATCCGGTTTGTT
TTTTGCGTTGAAAATACCCAGGTAATACAGGTTGTCGCGCATCAGAATGATCGCGTTGTTGGAATACTCTTTGG
ATTTAGACCAACCATCCGCCAGGGTCGGGATACCAAAGTTCAGTTTAATTTTTTTGGTGGAGTACGGCTTCTGA
GTTACGTAGTTACGTACCAGGTTATACAGAGAGATCACCGGATAGATTTCGTCATAGATTTCTTCCAGCTCAGC
ATAGAAGTTGTTATCTTTGTCAACCAGTTCTTCGGTCATAAACACAGAACACCAATGGAACGCATTCATAATGA
CATCCAGGACATTTTTCAGTTCAGAAGCCTTCAGTTCGCTTTCCACCAGGTGGATTTCCGGGTTATATTTCAGTT
CCTGGGCTTCGAAATTGTTCAAGATATGAGAAATTTCGTGAATGTATGTTTCCGCTTTGATGTTGTCATCAGAG
CACAGTTTGTAGTTAGACACCAGTTCGTTGATTTCGGTGATTGATTTTTGCAGGTCATTTTTTACCGCTTTTTTAA
CTTTATCGGCCTTGCTCTTGCCATTGCCCGGCAGGATGTTGTTGTAGTGGATTTCCAGTGCGGTGTTGATAGTTT
CCCAATCACGATAGGTCTTCTGAGAGACAGATTCATAGAATTTAGATACGATGTAGATTTTATCCAGATTATAA
CCATTGTAGTTATCGCCGATTTTGCGCAGACGTTCGACAATGTGTTTGCTGGAGATGTTATCTAGGAAACCGTT
TACGGACTGATACACCTCTTCGTCGGACTCAAATTTGTACGGTACTTCATAGCTAGTGTCCGCGATACAGAGGA
TCTGCTTATGCAGTTTCTGTAATTTGTACAGATTTTTGTTTTCTTTGTTTTTCTGACAGTACAGATTCATAAAACT
GTTCACTTTGCCACAAATATCGTTATAGAAACTGATACCTTCCTGGGTAATAAACTCACCGTATTTTTCGTAAGA
GTAGATTTCTTCCAGAGACATTTCTTTCAGTGAATCTTTCATGTCACCAGAGATTTTGTTAATGTCATCGTTGGA
AAGGCTTTTGACGATGCGACGGTAAACCAGAGCGTTGCTAAAGAAGATTTCCGCGTTGTCGTTAACAATGCGG
TGACAGGAGCTTGAGCTGATGTCATCAGCAGAGAAGCAGTTTGCACGGTTTTTAAAATAGTCCTTGAACGAAG
TAGCGAAACGGCTGAAAAGTTTAATTACCTGAGTTTTTTCTTCTTTCTCGGAGGCAGAGTAGTTGTTGTTGTGA
ATGACGAACTCCGGCAGAATATCGCTAATCAGCTTGGCGGAGAACATATTTTTGAAGCGATCATCGTTCGCGA
ACTTTTTATGAATGGCTTTACGATACTCTGTCTGTTCCTTGATCAGGGTATCTTTGTTATCACCGTTTTTTAACTG
GATTTCCATTTTTTCGAACAGACTGGTCCAATCGATGTCATCGATGCTGGACAGGGTTTCGCTGATAAAGCCAC
GGTAGTAATCGTCCATAATATCCTTCAGAATCTGACGGTTTTCCCCACGCAGCTCGTCCTCTTTGATAATACCGT
TTTTAACAATGAACTGTTGGGTCGTCTCGGTAGGGATGAGCGCGTTGCGCAGGGTTTTCTGGAGGGAAGAGA
TACCGATGAAGTTTTGGAAGTTGTTAGTGCCGTTATTCATTTTTTGCTTTCTCCTCTTTTCATCCAAAATACGCCA
TGAATATCTCCAACGAGATAACACGGTTAAATCCTTCACCGGGGGATCTGCTCAATATTAACTCTACCGATATC
TTCGGCTTATGCCGAGCACCCCTGGCTACTAGTATTATACCTAGGACTGAGCTAGCTGTCAACCGATCATTGTC
AGTTAGCCATGGATTCTTCGTCTGTTTCTACTGGTATTGGCACAAACCTGATTCCAATTTGAGCAAGGCTATGTG
CCATCTCGATACTCGTTCTTAACTCAACAGAAGATGCTTTGTGCATACAGCCCCTCGTTTATTATTTATCTCCTCA
GCCAGCCGCTGTGCTTTCAGTGGATTTCGGATAACAGAAAGGCCGGGAAATACCCAGCCTCGCTTTGTAACGG
AGTAGACGAAAGTGATTGCGCCTACCCGGATATTATCGTGAGGATGCGTCATCGCCATTGCTCCCCAAATACA
AAACCAATTTCAGCCAGTGCCTCGTCCATTTTTTCGATGAACTCCGGCACGATCTCGTCAAAACTCGCCATGTAC
TTTTCATCCCGCTCAATCACGACATAATGCAGGCCTTCACGCTTCATACGCGGGTCATAGTTGGCAAAGTACCA
GGCATTTTTTCGCGTCACCCACATGCTGTACTGCACCTGGGCCATGTAAGCTGACTTTATGGCCTCGAAACCAC
CGAGCCGGAACTTCATGAAATCCCGGGAGGTAAACGGGCATTTCAGTTCAAGGCCGTTGCCGTCACTGCATAA
ACCATCGGGAGAGCAGGCGGTACGCATACTTTCGTCGCGATAGATGATCGGGGATTCAGTAACATTCACGCCG
GAAGTGAATTCAAACAGGGTTCTGGCGTCGTTCTCGTACTGTTTTCCCCAGGCCAGTGCTTTAGCGTTAACTTC
CGGAGCCACACCGGTGCAAACCTCAGCAAGCAGGGTGTGGAAGTAGGACATTTTCATGTCAGGCCACTTCTTT
CCGGAGCGGGGTTTTGCTATCACGTTGTGAACTTCTGAAGCGGTGATGACGCCGAGCCGTAATTTGTGCCACG
CATCATCCCCCTGTTCGACAGCTCTCACATCGATCCCGGTACGCTGCAGGATAATGTCCGGTGTCATGCTGCCA
CCTTCTGCTCTGCGGCTTTCTGTTTCAGGAATCCAAGAGCTTTTACTGCTTCGGCCTGTGTCAGTTCTGACGATG
CACGAATGTCGCGGCGAAATATCTGGGAACAGAGCGGCAATAAGTCGTCATCCCATGTTTTATCCAGGGCGAT
CAGCAGAGTGTTAATCTCCTGCATGGTTTCATCGTTAACCGGAGTGATGTCGCGTTCCGGCTGACGTTCTGCAG
TGTATGCAGTATTTTCGACAATGCGCTCGGCTTCATCCTTGTCATAGATACCAGCAAATCCGAAGGCCAGACGG

GCACACTGAATCATGGCTTTATGACGTAACATCCGTTTGGGATGCGACTGCCACGGCCCCGTGATTTCTCTGCC
TTCGCGAGTTTTGAATGGTTCGCGGCGGCATTCATCCATCCATTCGGTAACGCAGATCGGATGATTACGGTCCT
TGCGGTAAATCCGGCATGTACAGGATTCATTGTCCTGCTCAAAGTCCATGCCATCAAACTGCTGGTTTTCATTG
ATGATGCGGGACCAGCCATCAACGCCCACCACCGGAACGATGCCATTCTGCTTATCAGGAAAGGCGTAAATTT
CTTTCGTCCACGGATTAAGGCCGTACTGGTTGGCAACGATCAGTAATGCGATGAACTGCGCATCGCTGGCATC
ACCTTTAAATGCCGTCTGGCGAAGAGTGGTGATCAGTTCCTGTGGGTCGACAGAATCCATGCCGACACGTTCA
GCCAGCTTCCCAGCCAGCGTTGCGAGTGCAGTACTCATTCGTTTTATACCTCTGAATCAATATCAACCTGGTGG
TGAGCAATGGTTTCAACCATGTACCGGATGTGTTCTGCCATGCGCTCCTGAAACTCAACATCGTCATCAAACGC
ACGGGTAATGGATTTTTTGCTGGCCCCGTGGCGTTGCAAATGATCGATGCATAGCGATTCAAACAGGTGCTGG
GGCAGGCCTTTTTCCATGTCGTCTGCCAGTTCTGCCTCTTTCTCTTCACGGGCGAGCTGCTGGTAGTGACGCGC
CCAGCTCTGAGCCTCAAGACGATCCTGAATGTAATAAGCGTTCATGGCTGAACTCCTGAAATAGCTGTGAAAA
TATCGCCCGCGAAATGCCGGGCTGATTAGGAAAACAGGAAAGGGGGTTAGTGAATGCTTTTGCTTGATCTCAG
TTTCAGTATTAATATCCATTTTTTATAACCTCCTTAGAGCTCGAATTCCCAAAAAAACGGGTATGGAGAAACAGT
AGAGAGTTGCGATAAAAAGCGTCAGGTAGGATCCGCTAATCTTATGGATAAAAATGCTATGGCATAGCAAAGT
GTGACGCCGTGCAAATAATCAATGTGGACTTTTCTGCCGTGATTATAGACACTTTTGTTACGCGTTTTTGTCATG
GCTTTGGTCCCGCTTTGTTACAGAATGCTTTTAATAAGCGGGGTTACCGGTTTGGTTAGCGAGAAGAGCCAGT
AAAAGACGCAGTGACGGCAATGTCTGATGCAATATGGACAATTGGTTTCTTCTCTGAATGGCGGGAGTATGAA
AAGTATGGCTGAAGCGCAAAATGATCCCCTGCTGCCGGGATACTCGTTTAATGCCCATCTGGTGGCGGGTTTA
ACGCCGATTGAGGCCAACGGTTATCTCGATTTTTTTATCGACCGACCGCTGGGAATGAAAGGTTATATTCTCAA
TCTCACCATTCGCGGTCAGGGGGTGGTGAAAAATCAGGGACGAGAATTTGTTTGCCGACCGGGTGATATTTTG
CTGTTCCCGCCAGGAGAGATTCATCACTACGGTCGTCATCCGGAGGCTCGCGAATGGTATCACCAGTGGGTTT
ACTTTCGTCCGCGCGCCTACTGGCATGAATGGCTTAACTGGCCGTCAATATTTGCCAATACGGGGTTCTTTCGC
CCGGATGAAGCGCACCAGCCGCATTTCAGCGACCTGTTTGGGCAAATCATTAACGCCGGGCAAGGGGAAGGG
CGCTATTCGGAGCTGCTGGCGATAAATCTGCTTGAGCAATTGTTACTGCGGCGCATGGAAGCGATTAACGAGT
CGCTCCATCCACCGATGGATAATCGGGTACGCGAGGCTTGTCAGTACATCAGCGATCACCTGGCAGACAGCAA
TTTTGATATCGCCAGCGTCGCACAGCATGTTTGCTTGTCGCCGTCGCGTCTGTCACATCTTTTCCGCCAGCAGTT
AGGGATTAGCGTCTTAAGCTGGCGCGAGGACCAACGTATCAGCCAGGCGAAGCTGCTTTTGAGCACCACCCG
GATGCCTATCGCCACCGTCGGTCGCAATGTTGGTTTTGACGATCAACTCTACTTCTCGCGGGTATTTAAAAAAT
GCACCGGGGCCAGCCCGAGCGAGTTCCGTGCCGGTTGTGAAGAAAAAGTGAATGATGTAGCCGTCAAGTTGT
CATAATAAATCGATGCAGGTGGCACTTTTCGGGGAAATGTGCGCGGAACCCCTATTTGTTTATTTTTCTAAATA
CATTCAAATATGTATCCGCTCATGAGACAATAACCCTGATAAATGCTTCAATAATATTGAAAAAGGAAGAGTAT
GAGTATTCAACATTTCCGTGTCGCCCTTATTCCCTTTTTTGCGGCATTTTGCCTTCCTGTTTTTGCTCACCCAGAA
ACGCTGGTGAAAGTAAAAGATGCTGAAGATCAGTTGGGTGCACGAGTGGGTTACATCGAACTGGATCTCAAC
AGCGGTAAGATCCTTGAGAGTTTTCGCCCCGAAGAACGTTTTCCAATGATGAGCACTTTTAAAGTTCTGCTATG
TGGCGCGGTATTATCCCGTATTGACGCCGGGCAAGAGCAACTCGGTCGCCGCATACACTATTCTCAGAATGAC
TTGGTTGAGTACTCACCAGTCACAGAAAAGCATCTTACGGATGGCATGACAGTAAGAGAATTATGCAGTGCTG
CCATAACCATGAGTGATAACACTGCGGCCAACTTACTTCTGACAACGATCGGAGGACCGAAGGAGCTAACCGC
TTTTTTGCACAACATGGGGGATCATGTAACTCGCCTTGATCGTTGGGAACCGGAGCTGAATGAAGCCATACCA
AACGACGAGCGTGACACCACGATGCCTGTAGCAATGGCAACAACGTTGCGCAAACTATTAACTGGCGAACTAC
TTACTCTAGCTTCCCGGCAACAATTAATAGACTGGATGGAGGCGGATAAAGTTGCAGGACCACTTCTGCGCTC
GGCCCTTCCGGCTGGCTGGTTTATTGCTGATAAATCTGGAGCCGGTGAGCGTGGGTCTCGCGGTATCATTGCA
GCACTGGGGCCAGATGGTAAGCCCTCCCGTATCGTAGTTATCTACACGACGGGGAGTCAGGCAACTATGGAT
GAACGAAATAGACAGATCGCTGAGATAGGTGCCTCACTGATTAAGCATTGGTAACTGTCAGACCAAGTTTACT
CATATATACTTTAGATTGATTTAAAACTTCATTTTTAATTTAAAAGGATCTAGGTGAAGATCCTTTTTGATAATCT
CATGACCAAAATCCCTTAACGTGAGTTTTCGTTCCACTGAGCGTCAGACCCC

**>pGRNA-galK**

TTGAGATCCTTTTTTTCTGCGCGTAATCTGCTGCTTGCAAACAAAAAAACCACCGCTACCAGCGGTGGTTTGTTT
GCCGGATCAAGAGCTACCAACTCTTTTTCCGAAGGTAACTGGCTTCAGCAGAGCGCAGATACCAAATACTGTTC
TTCTAGTGTAGCCGTAGTTAGGCCACCACTTCAAGAACTCTGTAGCACCGCCTACATACCTCGCTCTGCTAATCC
TGTTACCAGTGGCTGCTGCCAGTGGCGATAAGTCGTGTCTTACCGGGTTGGACTCAAGACGATAGTTACCGGA
TAAGGCGCAGCGGTCGGGCTGAACGGGGGGTTCGTGCACACAGCCCAGCTTGGAGCGAACGACCTACACCG
AACTGAGATACCTACAGCGTGAGCTATGAGAAAGCGCCACGCTTCCCGAAGGGAGAAAGGCGGACAGGTATC
CGGTAAGCGGCAGGGTCGGAACAGGAGAGCGCACGAGGGAGCTTCCAGGGGGAAACGCCTGGTATCTTTAT
AGTCCTGTCGGGTTTCGCCACCTCTGACTTGAGCGTCGATTTTTGTGATGCTCGTCAGGGGGGCGGAGCCTAT
GGAAAAACGCCAGCAACGCGGCCTTTTTACGGTTCCTGGCCTTTTGCTGGCCTTTTGCTCACATGTTCTTTCCTG
CGTTATCCCCTGATTCTGTGGATAACCGTATTACCGCCTTTGAGTGAGCTGATACCGCTCGCCGCAGCCGAACG
ACCGAGCGCAGCGAGTGCCATCGTATTAGCACCCCGCACTCTAGATTTCAGTGCAATTTATCTCTTCAAATGTA
GCACCTGAAGTCAGCCCCATACGATATAAGTTGTAATTCTCATGTTTGACAGCTTATCATCGATAAGCTTTAATG
CGGTAGTTTATCACAGTTAAATTGCTACCAATTTCTACTCTTGTAGATGCACTGTTCACGACGGGTGTTGTAATT
TCTACTCTTGTAGATAGCTGCCACATCAAACTGGAGAATAAGGTCAGTTTCACCTGTTTTACGTAAAAACCCGC
TTCGGCGGGTTTTTACTTTTGGCTCGAGCAGCTCGATGTTAGCGTCGGCGTCCCGGGAGCTGCATGTGTCAGA
GGTTTTCACCGTCATCACCGAAACGCGCGAGACGAAAGGGCCTCGTGATACGCCTATTTTTATAGGTTAATGTC
ATGATAATAATGGTTTCTTAGACGTCAGGTGGCACTTTTCGGGAAGTTCCTATTCTCTAGAAAGTATAGGAACT
TCGGAAATGTGCGCGGAACCCCTATTTGTTTATTTTTCTAAATACATTCAAATATGTATCCGCTCATGAGACAAT
AACCCTGATAAATGCTTCAATAATATTGAAAAAGGAAGAGTATGATTGAACAAGATGGATTGCACGCAGGTTC
TCCGGCCGCTTGGGTGGAGAGGCTATTCGGCTATGACTGGGCACAACAGACAATCGGCTGCTCTGATGCCGCC
GTGTTCCGGCTGTCAGCGCAGGGGCGCCTGGTTCTTTTTGTCAAGACCGACCTCTCCGGTGCCCTGAATGAACT
GCAAGACGAGGCAGCGCGGCTATCGTGGCTGGCCACGACGGGCGTTCCTTGCGCAGCTGTGCTCGACGTTGT
CACTGAAGCGGGAAGGGACTGGCTGCTATTGGGCGAAGTGCCGGGGCAGGATCTCCTGTCATCTCACCTTGCT
CCTGCCGAGAAAGTATCCATCATGGCTGATGCAATGCGGCGGCTGCATACGCTTGATCCGGCTACCTGCCCATT
CGACCACCAAGCGAAACATCGCATCGAGCGAGCACGTACTCGGATGGAAGCCGGTCTTGTCGATCAGGATGA
TCTGGACGAAGAGCATCAGGGGCTCGCGCCAGCCGAACTGTTCGCCAGGCTCAAGGCGAGCATGCCCGACGG
CGAGGATCTCGTCGTGACCCATGGCGATGCCTGCTTGCCGAATATCATGGTGGAAAATGGCCGCTTTTCTGGA
TTCATCGACTGTGGCCGGCTGGGTGTGGCGGACCGCTATCAGGACATAGCGTTGGCTACCCGTGATATTGCTG
AAGAGCTTGGCGGCGAATGGGCTGACCGCTTCCTCGTGCTTTACGGTATCGCCGCTCCCGATTCGCAGCGCAT
CGCCTTCTATCGCCTTCTTGACGAGTTCTTCTGACTGTCAGACCAAGTTTACTCATATATACTTTAGATTGATTTA
AAACTTCATTTTTAATTTAAAAGGATCTAGGTGAAGATCCTTTTTGATAATCTCATGACCAAAATCCCTTAACGT
GAGTTTTCGTTCCACTGAAGTTCCTATTCTCTAGAAAGTATAGGAACTTCGAGCGTCAGACCCCGTAGAAAAGA
TCAAAGGATCTTC

**>pGRNA-galK-Donor**

TTGAGATCCTTTTTTTCTGTGCGTAATCTGCTGCTTGCAAACAAAAAAACCACCGCTACCAGCGGTGGTTTGTTT
GCCGGATCAAGAGCTACCAACTCTTTTTCCGAAGGTAACTGGCTTCAGCAGAGCGCAGATACCAAATACTGTTC
TTCTAGTGTAGCCGTAGTTAGGCCACCACTTCAAGAACTCTGTAGCACCGCCTACATACCTCGCTCTGCTAATCC
TGTTACCAGTGGCTGCTGCCAGTGGCGATAAGTCGTGTCTTACCGGGTTGGACTCAAGACGATAGTTACCGGA
TAAGGCGCAGCGGTCGGGCTGAACGGGGGGTTCGTGCACACAGCCCAGCTTGGAGCGAACGACCTACACCG
AACTGAGATACCTACAGCGTGAGCTATGAGAAAGCGCCACGCTTCCCGAAGGGAGAAAGGCGGACAGGTATC
CGGTAAGCGGCAGGGTCGGAACAGGAGAGCGCACGAGGGAGCTTCCAGGGGGAAACGCCTGGTATCTTTAT
AGTCCTGTCGGGTTTCGCCACCTCTGACTTGAGCGTCGATTTTTGTGATGCTCGTCAGGGGGGCGGAGCCTAT
GGAAAAACGCCAGCAACGCGGCCTTTTTACGGTTCCTGGCCTTTTGCTGGCCTTTTGCTCACATGTTCTTTCCTG
CGTTATCCCCTGATTCTGAACAGTAACTTCAAACGTACCCTGGTTGGCAGCGAATACAACACCTAGGGTGAACA
GTGCGAGACCGGTGCGCGTTTCTTCCAGCAGCCAGCCCTGCGTGATGTTGAGTGAGCTGATACCGCTCGCCGC
AGCCGAACGACCGAGCGCAGCGAGTGCCATCGTATTAGCACCCCGCACTCTAGATTTCAGTGCAATTTATCTCT
TCAAATGTAGCACCTGAAGTCAGCCCCATACGATATAAGTTGTAATTCTCATGTTTGACAGCTTATCATCGATAA
GCTTTAATGCGGTAGTTTATCACAGTTAAATTGCTACCAATTTCTACTCTTGTAGATGCACTGTTCACGACGGGT
GTTGTAATTTCTACTCTTGTAGATAGCTGCCACATCAAACTGGAGAATAAGGTCAGTTTCACCTGTTTTACGTAA
AAACCCGCTTCGGCGGGTTTTTACTTTTGGCTCGAGCAGCTCGATGTTAGCGTCGGCGTCCCGGGAGCTGCAT
GTGTCAGAGGTTTTCACCGTCATCACCGAAACGCGCGAGACGAAAGGGCCTCGTGATACGCCTATTTTTATAG
GTTAATGTCATGATAATAATGGTTTCTTAGACGTCAGGTGGCACTTTTCGGGAAGTTCCTATTCTCTAGAAAGT
ATAGGAACTTCGGAAATGTGCGCGGAACCCCTATTTGTTTATTTTTCTAAATACATTCAAATATGTATCCGCTCA
TGAGACAATAACCCTGATAAATGCTTCAATAATATTGAAAAAGGAAGAGTATGATTGAACAAGATGGATTGCA
CGCAGGTTCTCCGGCCGCTTGGGTGGAGAGGCTATTCGGCTATGACTGGGCACAACAGACAATCGGCTGCTCT
GATGCCGCCGTGTTCCGGCTGTCAGCGCAGGGGCGCCTGGTTCTTTTTGTCAAGACCGACCTCTCCGGTGCCCT
GAATGAACTGCAAGACGAGGCAGCGCGGCTATCGTGGCTGGCCACGACGGGCGTTCCTTGCGCAGCTGTGCT
CGACGTTGTCACTGAAGCGGGAAGGGACTGGCTGCTATTGGGCGAAGTGCCGGGGCAGGATCTCCTGTCATC
TCACCTTGCTCCTGCCGAGAAAGTATCCATCATGGCTGATGCAATGCGGCGGCTGCATACGCTTGATCCGGCTA
CCTGCCCATTCGACCACCAAGCGAAACATCGCATCGAGCGAGCACGTACTCGGATGGAAGCCGGTCTTGTCGA
TCAGGATGATCTGGACGAAGAGCATCAGGGGCTCGCGCCAGCCGAACTGTTCGCCAGGCTCAAGGCGAGCAT
GCCCGACGGCGAGGATCTCGTCGTGACCCATGGCGATGCCTGCTTGCCGAATATCATGGTGGAAAATGGCCG
CTTTTCTGGATTCATCGACTGTGGCCGGCTGGGTGTGGCGGACCGCTATCAGGACATAGCGTTGGCTACCCGT
GATATTGCTGAAGAGCTTGGCGGCGAATGGGCTGACCGCTTCCTCGTGCTTTACGGTATCGCCGCTCCCGATTC
GCAGCGCATCGCCTTCTATCGCCTTCTTGACGAGTTCTTCTGACTGTCAGACCAAGTTTACTCATATATACTTTA
GATTGATTTAAAACTTCATTTTTAATTTAAAAGGATCTAGGTGAAGATCCTTTTTGATAATCTCATGACCAAAAT
CCCTTAACGTGAGTTTTCGTTCCACTGAAGTTCCTATTCTCTAGAAAGTATAGGAACTTCGAGCGTCAGACCCC
GTAGAAAAGATCAAAGGATCTTC

**>pGRNA-sacB-galK**

TTGAGATCCTTTTTTTCTGCGCGTAATCTGCTGCTTGCAAACAAAAAAACCACCGCTACCAGCGGTGGTTTGTTT
GCCGGATCAAGAGCTACCAACTCTTTTTCCGAAGGTAACTGGCTTCAGCAGAGCGCAGATACCAAATACTGTTC
TTCTAGTGTAGCCGTAGTTAGGCCACCACTTCAAGAACTCTGTAGCACCGCCTACATACCTCGCTCTGCTAATCC
TGTTACCAGTGGCTGCTGCCAGTGGCGATAAGTCGTGTCTTACCGGGTTGGACTCAAGACGATAGTTACCGGA
TAAGGCGCAGCGGTCGGGCTGAACGGGGGGTTCGTGCACACAGCCCAGCTTGGAGCGAACGACCTACACCG
AACTGAGATACCTACAGCGTGAGCTATGAGAAAGCGCCACGCTTCCCGAAGGGAGAAAGGCGGACAGGTATC
CGGTAAGCGGCAGGGTCGGAACAGGAGAGCGCACGAGGGAGCTTCCAGGGGGAAACGCCTGGTATCTTTAT
AGTCCTGTCGGGTTTCGCCACCTCTGACTTGAGCGTCGATTTTTGTGATGCTCGTCAGGGGGGCGGAGCCTAT
GGAAAAACGCCAGCAACGCGGCCTTTTTACGGTTCCTGGCCTTTTGCTGGCCTTTTGCTCACATGTTCAAGCTTA
GAGGTTCCAACTTTCACCATAATGAAACACTCGAGAATAATTTTGTTTAACTTTAAGAAGGAGATATACATATG
AACATTAAAAAATTTGCGAAACAGGCGACAGTGCTGACCTTTACAACCGCGCTGTTAGCGGGCGGCGCGACCC
AGGCGTTTGCGAAAGAAACCAACCAGAAACCGTATAAAGAAACCTATGGCATTAGCCATATTACCCGCCATGA
TATGCTGCAGATCCCGGAACAGCAGAAAAACGAAAAATATCAGGTGCCGGAATTTGATAGCAGCACCATTAAA
AACATTAGCAGCGCGAAAGGCCTGGATGTGTGGGATAGCTGGCCGCTGCAGAACGCGGATGGCACCGTGGC
GAACTATCATGGCTATCATATTGTGTTTGCGCTGGCGGGCGATCCGAAAAACGCGGATGATACCAGTATTTAT
ATGTTTTATCAGAAAGTGGGCGAAACCAGCATTGATAGCTGGAAAAACGCGGGCCGCGTGTTTAAAGATAGC
GATAAATTTGATGCGAACGATAGCATTCTGAAAGATCAGACCCAGGAATGGTCAGGCAGCGCGACCTTTACCA
GCGATGGCAAAATTCGCCTGTTTTATACCGATTTTAGCGGCAAACATTATGGCAAACAGACCCTGACCACCGCC
CAGGTGAACGTGAGCGCGAGCGATAGCAGCCTGAACATTAACGGCGTGGAGGACTACAAGAGCATTTTTGAT
GGCGATGGCAAAACCTATCAGAACGTGCAACAGTTTATTGATGAAGGCAACTATAGCAGCGGCGATAACCATA
CCCTGCGCGATCCGCATTATGTGGAAGATAAAGGCCATAAATATCTGGTGTTTGAAGCGAACACCGGCACCGA
AGATGGCTATCAGGGCGAAGAAAGCCTGTTTAACAAAGCGTATTATGGCAAAAGCACCAGCTTTTTTCGCCAG
GAAAGCCAGAAACTGCTGCAGAGCGATAAAAAACGCACCGCGGAACTGGCGAACGGTGCGCTGGGCATGAT
TGAACTGAACGATGATTATACCCTGAAAAAAGTGATGAAACCGCTGATTGCGAGCAACACAGTGACAGATGA
AATTGAACGCGCGAATGTTTTTAAAATGAACGGCAAATGGTATCTGTTTACCGATAGCCGCGGCAGCAAAATG
ACCATTGATGGCATTACCAGCAACGATATTTATATGCTGGGCTATGTGAGCAACAGTCTGACAGGCCCGTATA
AACCGCTGAACAAAACCGGCCTGGTGCTGAAAATGGATTTAGATCCGAACGATGTGACCTTTACCTATAGCCA
TTTTGCGGTGCCGCAGGCGAAAGGCAACAACGTGGTGATTACCAGCTATATGACCAACCGCGGCTTTTATGCG
GATAAACAGAGTACCTTTGCGCCGAGCTTTCTGCTGAACATTAAAGGCAAAAAAACCAGCGTGGTGAAAGATA
GCATTCTGGAACAGGGCCAACTTACCGTGAACAAAGGTACCTAATAGCTGAAGATCCGGCTGCTAACAAAGCC
CGAAAGGAAGCTGAGTTGGCTGCTGCCACCGCTGAGCAATAACTAGCATAACCCCTTGGGGCCTCTAAACGG
GTCTTGAGGGGTTTTTTGCTGAAAGGATCCACGTCAGGTTGCCATCGTATTAGCACCCCGCACTCTAGATTTCA
GTGCAATTTATCTCTTCAAATGTAGCACCTGAAGTCAGCCCCATACGATATAAGTTGTAATTCTCATGTTTGACA
GCTTATCATCGATAAGCTTTAATGCGGTAGTTTATCACAGTTAAATTGCTACCAATTTCTACTCTTGTAGATGCA
CTGTTCACGACGGGTGTTGTAATTTCTACTCTTGTAGATAGCTGCCACATCAAACTGGAGAATAAGGTCAGTTT
CACCTGTTTTACGTAAAAACCCGCTTCGGCGGGTTTTTACTTTTGGCTCGAGCAGCTCGATGTTAGCGTCGGCG
TCCCGGGAGCTGCATGTGTCAGAGGTTTTCACCGTCATCACCGAAACGCGCGAGACGAAAGGGCCTCGTGATA
CGCCTATTTTTATAGGTTAATGTCATGATAATAATGGTTTCTTAGACGTCAGGTGGCACTTTTCGGGAAGTTCCT
ATTCTCTAGAAAGTATAGGAACTTCGGAAATGTGCGCGGAACCCCTATTTGTTTATTTTTCTAAATACATTCAAA
TATGTATCCGCTCATGAGACAATAACCCTGATAAATGCTTCAATAATATTGAAAAAGGAAGAGTATGATTGAAC
AAGATGGATTGCACGCAGGTTCTCCGGCCGCTTGGGTGGAGAGGCTATTCGGCTATGACTGGGCACAACAGA
CAATCGGCTGCTCTGATGCCGCCGTGTTCCGGCTGTCAGCGCAGGGGCGCCTGGTTCTTTTTGTCAAGACCGA
CCTCTCCGGTGCCCTGAATGAACTGCAAGACGAGGCAGCGCGGCTATCGTGGCTGGCCACGACGGGCGTTCCT
TGCGCAGCTGTGCTCGACGTTGTCACTGAAGCGGGAAGGGACTGGCTGCTATTGGGCGAAGTGCCGGGGCAG
GATCTCCTGTCATCTCACCTTGCTCCTGCCGAGAAAGTATCCATCATGGCTGATGCAATGCGGCGGCTGCATAC
GCTTGATCCGGCTACCTGCCCATTCGACCACCAAGCGAAACATCGCATCGAGCGAGCACGTACTCGGATGGAA
GCCGGTCTTGTCGATCAGGATGATCTGGACGAAGAGCATCAGGGGCTCGCGCCAGCCGAACTGTTCGCCAGG

CTCAAGGCGAGCATGCCCGACGGCGAGGATCTCGTCGTGACCCATGGCGATGCCTGCTTGCCGAATATCATGG
TGGAAAATGGCCGCTTTTCTGGATTCATCGACTGTGGCCGGCTGGGTGTGGCGGACCGCTATCAGGACATAGC
GTTGGCTACCCGTGATATTGCTGAAGAGCTTGGCGGCGAATGGGCTGACCGCTTCCTCGTGCTTTACGGTATC
GCCGCTCCCGATTCGCAGCGCATCGCCTTCTATCGCCTTCTTGACGAGTTCTTCTGACTGTCAGACCAAGTTTAC
TCATATATACTTTAGATTGATTTAAAACTTCATTTTTAATTTAAAAGGATCTAGGTGAAGATCCTTTTTGATAATC
TCATGACCAAAATCCCTTAACGTGAGTTTTCGTTCCACTGAAGTTCCTATTCTCTAGAAAGTATAGGAACTTCGA
GCGTCAGACCCCGTAGAAAAGATCAAAGGATCTTC

**>pGRNA-sacB-ccdB**

TTGAGATCCTTTTTTTCTGTGCGTAATCTGCTGCTTGCAAACAAAAAAACCACCGCTACCAGCGGTGGTTTGTTT
GCCGGATCAAGAGCTACCAACTCTTTTTCCGAAGGTAACTGGCTTCAGCAGAGCGCAGATACCAAATACTGTTC
TTCTAGTGTAGCCGTAGTTAGGCCACCACTTCAAGAACTCTGTAGCACCGCCTACATACCTCGCTCTGCTAATCC
TGTTACCAGTGGCTGCTGCCAGTGGCGATAAGTCGTGTCTTACCGGGTTGGACTCAAGACGATAGTTACCGGA
TAAGGCGCAGCGGTCGGGCTGAACGGGGGGTTCGTGCACACAGCCCAGCTTGGAGCGAACGACCTACACCG
AACTGAGATACCTACAGCGTGAGCTATGAGAAAGCGCCACGCTTCCCGAAGGGAGAAAGGCGGACAGGTATC
CGGTAAGCGGCAGGGTCGGAACAGGAGAGCGCACGAGGGAGCTTCCAGGGGGAAACGCCTGGTATCTTTAT
AGTCCTGTCGGGTTTCGCCACCTCTGACTTGAGCGTCGATTTTTGTGATGCTCGTCAGGGGGGCGGAGCCTAT
GGAAAAACGCCAGCAACGCGGCCTTTTTACGGTTCCTGGCCTTTTGCTGGCCTTTTGCTCACATGTTCAAGCTTA
GAGGTTCCAACTTTCACCATAATGAAACACTCGAGAATAATTTTGTTTAACTTTAAGAAGGAGATATACATATG
AACATTAAAAAATTTGCGAAACAGGCGACAGTGCTGACCTTTACAACCGCGCTGTTAGCGGGCGGCGCGACCC
AGGCGTTTGCGAAAGAAACCAACCAGAAACCGTATAAAGAAACCTATGGCATTAGCCATATTACCCGCCATGA
TATGCTGCAGATCCCGGAACAGCAGAAAAACGAAAAATATCAGGTGCCGGAATTTGATAGCAGCACCATTAAA
AACATTAGCAGCGCGAAAGGCCTGGATGTGTGGGATAGCTGGCCGCTGCAGAACGCGGATGGCACCGTGGC
GAACTATCATGGCTATCATATTGTGTTTGCGCTGGCGGGCGATCCGAAAAACGCGGATGATACCAGTATTTAT
ATGTTTTATCAGAAAGTGGGCGAAACCAGCATTGATAGCTGGAAAAACGCGGGCCGCGTGTTTAAAGATAGC
GATAAATTTGATGCGAACGATAGCATTCTGAAAGATCAGACCCAGGAATGGTCAGGCAGCGCGACCTTTACCA
GCGATGGCAAAATTCGCCTGTTTTATACCGATTTTAGCGGCAAACATTATGGCAAACAGACCCTGACCACCGCC
CAGGTGAACGTGAGCGCGAGCGATAGCAGCCTGAACATTAACGGCGTGGAGGACTACAAGAGCATTTTTGAT
GGCGATGGCAAAACCTATCAGAACGTGCAACAGTTTATTGATGAAGGCAACTATAGCAGCGGCGATAACCATA
CCCTGCGCGATCCGCATTATGTGGAAGATAAAGGCCATAAATATCTGGTGTTTGAAGCGAACACCGGCACCGA
AGATGGCTATCAGGGCGAAGAAAGCCTGTTTAACAAAGCGTATTATGGCAAAAGCACCAGCTTTTTCGCCAG
GAAAGCCAGAAACTGCTGCAGAGCGATAAAAAACGCACCGCGGAACTGGCGAACGGTGCGCTGGGCATGAT
TGAACTGAACGATGATTATACCCTGAAAAAAGTGATGAAACCGCTGATTGCGAGCAACACAGTGACAGATGA
AATTGAACGCGCGAATGTTTTTAAAATGAACGGCAAATGGTATCTGTTTACCGATAGCCGCGGCAGCAAAATG
ACCATTGATGGCATTACCAGCAACGATATTTATATGCTGGGCTATGTGAGCAACAGTCTGACAGGCCCGTATA
AACCGCTGAACAAAACCGGCCTGGTGCTGAAAATGGATTTAGATCCGAACGATGTGACCTTTACCTATAGCCA
TTTTGCGGTGCCGCAGGCGAAAGGCAACAACGTGGTGATTACCAGCTATATGACCAACCGCGGCTTTTATGCG
GATAAACAGAGTACCTTTGCGCCGAGCTTTCTGCTGAACATTAAAGGCAAAAAAACCAGCGTGGTGAAAGATA
GCATTCTGGAACAGGGCCAACTTACCGTGAACAAAGGTACCTAATAGCTGAAGATCCGGCTGCTAACAAAGCC
CGAAAGGAAGCTGAGTTGGCTGCTGCCACCGCTGAGCAATAACTAGCATAACCCCTTGGGGCCTCTAAACGG
GTCTTGAGGGGTTTTTTGCTGAAAGGATCCACGTCAGGTTGCCATCGTATTAGCACCCCGCACTCTAGATTTCA
GTGCAATTTATCTCTTCAAATGTAGCACCTGAAGTCAGCCCCATACGATATAAGTTGTAATTCTCATGTTTGACA
GCTTATCATCGATAAGCTTTAATGCGGTAGTTTATCACAGTTAAATTGCTACCAATTTCTACTCTTGTAGATGGA
GACCACCCGAAGTATGTCAAAAAGAGGTATGCTATGAAGCAGCGTATTACAGTGACAGTTGACAGCGACAGC
TATCAGTTGCTCAAGGCATATATGATGTCAATATCTCCGGTCTGGTAAGCACAACCATGCAGAATGAAGCCCGT
CGTCTGCGTGCCGAACGCTGGAAAGCGGAAAATCAGGAAGGGATGGCTGAGGTCGCCCGGTTTATTGAAATG
AACGGCTCTTTTGCTGACGAGAACAGGGGCTGGTGAAATGCAGTTTAAGGTTTACACCTATAAAAGAGAGAG
CCGTTATCGTCTGTTTGTGGATGTACAGAGTGATATTATTGACACGCCCGGGCGACGGATGGTGATCCCCCTG
GCCAGTGCACGTCTGCTGTCAGATAAAGTCTCCCGTGAACTTTACCCGGTGGTGCATATCGGGGATGAAAGCT
GGCGCATGATGACCACCGATATGGCCAGTGTGCCGGTTAGCGTTATCGGGGAAGAAGTGGCTGATCTCAGCC
ACCGCGAAAATGACATCAAAAACGCCATTAACCTGATGTTCTGGGGAATATAAATGTCAGGCTCCCTTATACAC
AGCCAGTCTGCAGGTCTCGAATTTCTACTCTTGTAGATAGCTGCCACATCAAACTGGAGAATAAGGTCAGTTTC
ACCTGTTTTACGTAAAAACCCGCTTCGGCGGGTTTTTACTTTTGGCTCGAGCAGCTCGATGTTAGCGTCGGCGT
CCCGGGAGCTGCATGTGTCAGAGGTTTTCACCGTCATCACCGAAACGCGCGAGACGAAAGGGCCTCGTGATA
CGCCTATTTTTATAGGTTAATGTCATGATAATAATGGTTTCTTAGACGTCAGGTGGCACTTTTCGGGAAGTTCCT
ATTCTCTAGAAAGTATAGGAACTTCGGAAATGTGCGCGGAACCCCTATTTGTTTATTTTTCTAAATACATTCAAA

TATGTATCCGCTCATGAGACAATAACCCTGATAAATGCTTCAATAATATTGAAAAAGGAAGAGTATGATTGAAC
AAGATGGATTGCACGCAGGTTCTCCGGCCGCTTGGGTGGAGAGGCTATTCGGCTATGACTGGGCACAACAGA
CAATCGGCTGCTCTGATGCCGCCGTGTTCCGGCTGTCAGCGCAGGGGCGCCTGGTTCTTTTTGTCAAGACCGA
CCTCTCCGGTGCCCTGAATGAACTGCAAGACGAGGCAGCGCGGCTATCGTGGCTGGCCACGACGGGCGTTCCT
TGCGCAGCTGTGCTCGACGTTGTCACTGAAGCGGGAAGGGACTGGCTGCTATTGGGCGAAGTGCCGGGGCAG
GATCTCCTGTCATCTCACCTTGCTCCTGCCGAGAAAGTATCCATCATGGCTGATGCAATGCGGCGGCTGCATAC
GCTTGATCCGGCTACCTGCCCATTCGACCACCAAGCGAAACATCGCATCGAGCGAGCACGTACTCGGATGGAA
GCCGGTCTTGTCGATCAGGATGATCTGGACGAAGAGCATCAGGGGCTCGCGCCAGCCGAACTGTTCGCCAGG
CTCAAGGCGAGCATGCCCGACGGCGAGGATCTCGTCGTGACCCATGGCGATGCCTGCTTGCCGAATATCATGG
TGGAAAATGGCCGCTTTTCTGGATTCATCGACTGTGGCCGGCTGGGTGTGGCGGACCGCTATCAGGACATAGC
GTTGGCTACCCGTGATATTGCTGAAGAGCTTGGCGGCGAATGGGCTGACCGCTTCCTCGTGCTTTACGGTATC
GCCGCTCCCGATTCGCAGCGCATCGCCTTCTATCGCCTTCTTGACGAGTTCTTCTGACTGTCAGACCAAGTTTAC
TCATATATACTTTAGATTGATTTAAAACTTCATTTTTAATTTAAAAGGATCTAGGTGAAGATCCTTTTTGATAATC
TCATGACCAAAATCCCTTAACGTGAGTTTTCGTTCCACTGAAGTTCCTATTCTCTAGAAAGTATAGGAACTTCGA
GCGTCAGACCCCGTAGAAAAGATCAAAGGATCTTC

**>pGRNA-sacB-endA**

TTGAGATCCTTTTTTTCTGCGCGTAATCTGCTGCTTGCAAACAAAAAAACCACCGCTACCAGCGGTGGTTTGTTT
GCCGGATCAAGAGCTACCAACTCTTTTTCCGAAGGTAACTGGCTTCAGCAGAGCGCAGATACCAAATACTGTTC
TTCTAGTGTAGCCGTAGTTAGGCCACCACTTCAAGAACTCTGTAGCACCGCCTACATACCTCGCTCTGCTAATCC
TGTTACCAGTGGCTGCTGCCAGTGGCGATAAGTCGTGTCTTACCGGGTTGGACTCAAGACGATAGTTACCGGA
TAAGGCGCAGCGGTCGGGCTGAACGGGGGGTTCGTGCACACAGCCCAGCTTGGAGCGAACGACCTACACCG
AACTGAGATACCTACAGCGTGAGCTATGAGAAAGCGCCACGCTTCCCGAAGGGAGAAAGGCGGACAGGTATC
CGGTAAGCGGCAGGGTCGGAACAGGAGAGCGCACGAGGGAGCTTCCAGGGGGAAACGCCTGGTATCTTTAT
AGTCCTGTCGGGTTTCGCCACCTCTGACTTGAGCGTCGATTTTTGTGATGCTCGTCAGGGGGGCGGAGCCTAT
GGAAAAACGCCAGCAACGCGGCCTTTTTACGGTTCCTGGCCTTTTGCTGGCCTTTTGCTCACATGTTCAAGCTTA
GAGGTTCCAACTTTCACCATAATGAAACACTCGAGAATAATTTTGTTTAACTTTAAGAAGGAGATATACATATG
AACATTAAAAAATTTGCGAAACAGGCGACAGTGCTGACCTTTACAACCGCGCTGTTAGCGGGCGGCGCGACCC
AGGCGTTTGCGAAAGAAACCAACCAGAAACCGTATAAAGAAACCTATGGCATTAGCCATATTACCCGCCATGA
TATGCTGCAGATCCCGGAACAGCAGAAAAACGAAAAATATCAGGTGCCGGAATTTGATAGCAGCACCATTAAA
AACATTAGCAGCGCGAAAGGCCTGGATGTGTGGGATAGCTGGCCGCTGCAGAACGCGGATGGCACCGTGGC
GAACTATCATGGCTATCATATTGTGTTTGCGCTGGCGGGCGATCCGAAAAACGCGGATGATACCAGTATTTAT
ATGTTTTATCAGAAAGTGGGCGAAACCAGCATTGATAGCTGGAAAAACGCGGGCCGCGTGTTTAAAGATAGC
GATAAATTTGATGCGAACGATAGCATTCTGAAAGATCAGACCCAGGAATGGTCAGGCAGCGCGACCTTTACCA
GCGATGGCAAAATTCGCCTGTTTTATACCGATTTTAGCGGCAAACATTATGGCAAACAGACCCTGACCACCGCC
CAGGTGAACGTGAGCGCGAGCGATAGCAGCCTGAACATTAACGGCGTGGAGGACTACAAGAGCATTTTTGAT
GGCGATGGCAAAACCTATCAGAACGTGCAACAGTTTATTGATGAAGGCAACTATAGCAGCGGCGATAACCATA
CCCTGCGCGATCCGCATTATGTGGAAGATAAAGGCCATAAATATCTGGTGTTTGAAGCGAACACCGGCACCGA
AGATGGCTATCAGGGCGAAGAAAGCCTGTTTAACAAAGCGTATTATGGCAAAAGCACCAGCTTTTTTCGCCAG
GAAAGCCAGAAACTGCTGCAGAGCGATAAAAAACGCACCGCGGAACTGGCGAACGGTGCGCTGGGCATGAT
TGAACTGAACGATGATTATACCCTGAAAAAAGTGATGAAACCGCTGATTGCGAGCAACACAGTGACAGATGA
AATTGAACGCGCGAATGTTTTTAAAATGAACGGCAAATGGTATCTGTTTACCGATAGCCGCGGCAGCAAAATG
ACCATTGATGGCATTACCAGCAACGATATTTATATGCTGGGCTATGTGAGCAACAGTCTGACAGGCCCGTATA
AACCGCTGAACAAAACCGGCCTGGTGCTGAAAATGGATTTAGATCCGAACGATGTGACCTTTACCTATAGCCA
TTTTGCGGTGCCGCAGGCGAAAGGCAACAACGTGGTGATTACCAGCTATATGACCAACCGCGGCTTTTATGCG
GATAAACAGAGTACCTTTGCGCCGAGCTTTCTGCTGAACATTAAAGGCAAAAAAACCAGCGTGGTGAAAGATA
GCATTCTGGAACAGGGCCAACTTACCGTGAACAAAGGTACCTAATAGCTGAAGATCCGGCTGCTAACAAAGCC
CGAAAGGAAGCTGAGTTGGCTGCTGCCACCGCTGAGCAATAACTAGCATAACCCCTTGGGGCCTCTAAACGG
GTCTTGAGGGGTTTTTTGCTGAAAGGATCCACGTCAGGTTGCCATCGTATTAGCACCCCGCACTCTAGATTTCA
GTGCAATTTATCTCTTCAAATGTAGCACCTGAAGTCAGCCCCATACGATATAAGTTGTAATTCTCATGTTTGACA
GCTTATCATCGATAAGCTTTAATGCGGTAGTTTATCACAGTTAAATTGCTACCAATTTCTACTCTTGTAGATCATT
TTTGCGCACCTGATAGCCGAATTTCTACTCTTGTAGATCATTTTTGCGCACCTGATAGCCGAATTTCTACTCTTGT
AGATAGCTGCCACATCAAACTGGAGAATAAGGTCAGTTTCACCTGTTTTACGTAAAAACCCGCTTCGGCGGGTT
TTTACTTTTGGCTCGAGCAGCTCGATGTTAGCGTCGGCGTCCCGGGAGCTGCATGTGTCAGAGGTTTTCACCGT
CATCACCGAAACGCGCGAGACGAAAGGGCCTCGTGATACGCCTATTTTTATAGGTTAATGTCATGATAATAAT
GGTTTCTTAGACGTCAGGTGGCACTTTTCGGGAAGTTCCTATTCTCTAGAAAGTATAGGAACTTCGGAAATGTG
CGCGGAACCCCTATTTGTTTATTTTTCTAAATACATTCAAATATGTATCCGCTCATGAGACAATAACCCTGATAA
ATGCTTCAATAATATTGAAAAAGGAAGAGTATGATTGAACAAGATGGATTGCACGCAGGTTCTCCGGCCGCTT
GGGTGGAGAGGCTATTCGGCTATGACTGGGCACAACAGACAATCGGCTGCTCTGATGCCGCCGTGTTCCGGCT
GTCAGCGCAGGGGCGCCTGGTTCTTTTTGTCAAGACCGACCTCTCCGGTGCCCTGAATGAACTGCAAGACGAG
GCAGCGCGGCTATCGTGGCTGGCCACGACGGGCGTTCCTTGCGCAGCTGTGCTCGACGTTGTCACTGAAGCG
GGAAGGGACTGGCTGCTATTGGGCGAAGTGCCGGGGCAGGATCTCCTGTCATCTCACCTTGCTCCTGCCGAGA
AAGTATCCATCATGGCTGATGCAATGCGGCGGCTGCATACGCTTGATCCGGCTACCTGCCCATTCGACCACCAA
GCGAAACATCGCATCGAGCGAGCACGTACTCGGATGGAAGCCGGTCTTGTCGATCAGGATGATCTGGACGAA

GAGCATCAGGGGCTCGCGCCAGCCGAACTGTTCGCCAGGCTCAAGGCGAGCATGCCCGACGGCGAGGATCTC
GTCGTGACCCATGGCGATGCCTGCTTGCCGAATATCATGGTGGAAAATGGCCGCTTTTCTGGATTCATCGACTG
TGGCCGGCTGGGTGTGGCGGACCGCTATCAGGACATAGCGTTGGCTACCCGTGATATTGCTGAAGAGCTTGG
CGGCGAATGGGCTGACCGCTTCCTCGTGCTTTACGGTATCGCCGCTCCCGATTCGCAGCGCATCGCCTTCTATC
GCCTTCTTGACGAGTTCTTCTGACTGTCAGACCAAGTTTACTCATATATACTTTAGATTGATTTAAAACTTCATTT
TTAATTTAAAAGGATCTAGGTGAAGATCCTTTTTGATAATCTCATGACCAAAATCCCTTAACGTGAGTTTTCGTT
CCACTGAAGTTCCTATTCTCTAGAAAGTATAGGAACTTCGAGCGTCAGACCCCGTAGAAAAGATCAAAGGATC
TTC

**>pGRNA-sacB-fruR**

TTGAGATCCTTTTTTTCTGCGCGTAATCTGCTGCTTGCAAACAAAAAAACCACCGCTACCAGCGGTGGTTTGTTT
GCCGGATCAAGAGCTACCAACTCTTTTTCCGAAGGTAACTGGCTTCAGCAGAGCGCAGATACCAAATACTGTTC
TTCTAGTGTAGCCGTAGTTAGGCCACCACTTCAAGAACTCTGTAGCACCGCCTACATACCTCGCTCTGCTAATCC
TGTTACCAGTGGCTGCTGCCAGTGGCGATAAGTCGTGTCTTACCGGGTTGGACTCAAGACGATAGTTACCGGA
TAAGGCGCAGCGGTCGGGCTGAACGGGGGGTTCGTGCACACAGCCCAGCTTGGAGCGAACGACCTACACCG
AACTGAGATACCTACAGCGTGAGCTATGAGAAAGCGCCACGCTTCCCGAAGGGAGAAAGGCGGACAGGTATC
CGGTAAGCGGCAGGGTCGGAACAGGAGAGCGCACGAGGGAGCTTCCAGGGGGAAACGCCTGGTATCTTTAT
AGTCCTGTCGGGTTTCGCCACCTCTGACTTGAGCGTCGATTTTTGTGATGCTCGTCAGGGGGGCGGAGCCTAT
GGAAAAACGCCAGCAACGCGGCCTTTTTACGGTTCCTGGCCTTTTGCTGGCCTTTTGCTCACATGTTCAAGCTTA
GAGGTTCCAACTTTCACCATAATGAAACACTCGAGAATAATTTTGTTTAACTTTAAGAAGGAGATATACATATG
AACATTAAAAAATTTGCGAAACAGGCGACAGTGCTGACCTTTACAACCGCGCTGTTAGCGGGCGGCGCGACCC
AGGCGTTTGCGAAAGAAACCAACCAGAAACCGTATAAAGAAACCTATGGCATTAGCCATATTACCCGCCATGA
TATGCTGCAGATCCCGGAACAGCAGAAAAACGAAAAATATCAGGTGCCGGAATTTGATAGCAGCACCATTAAA
AACATTAGCAGCGCGAAAGGCCTGGATGTGTGGGATAGCTGGCCGCTGCAGAACGCGGATGGCACCGTGGC
GAACTATCATGGCTATCATATTGTGTTTGCGCTGGCGGGCGATCCGAAAAACGCGGATGATACCAGTATTTAT
ATGTTTTATCAGAAAGTGGGCGAAACCAGCATTGATAGCTGGAAAAACGCGGGCCGCGTGTTTAAAGATAGC
GATAAATTTGATGCGAACGATAGCATTCTGAAAGATCAGACCCAGGAATGGTCAGGCAGCGCGACCTTTACCA
GCGATGGCAAAATTCGCCTGTTTTATACCGATTTTAGCGGCAAACATTATGGCAAACAGACCCTGACCACCGCC
CAGGTGAACGTGAGCGCGAGCGATAGCAGCCTGAACATTAACGGCGTGGAGGACTACAAGAGCATTTTTGAT
GGCGATGGCAAAACCTATCAGAACGTGCAACAGTTTATTGATGAAGGCAACTATAGCAGCGGCGATAACCATA
CCCTGCGCGATCCGCATTATGTGGAAGATAAAGGCCATAAATATCTGGTGTTTGAAGCGAACACCGGCACCGA
AGATGGCTATCAGGGCGAAGAAAGCCTGTTTAACAAAGCGTATTATGGCAAAAGCACCAGCTTTTTTCGCCAG
GAAAGCCAGAAACTGCTGCAGAGCGATAAAAAACGCACCGCGGAACTGGCGAACGGTGCGCTGGGCATGAT
TGAACTGAACGATGATTATACCCTGAAAAAAGTGATGAAACCGCTGATTGCGAGCAACACAGTGACAGATGA
AATTGAACGCGCGAATGTTTTTAAAATGAACGGCAAATGGTATCTGTTTACCGATAGCCGCGGCAGCAAAATG
ACCATTGATGGCATTACCAGCAACGATATTTATATGCTGGGCTATGTGAGCAACAGTCTGACAGGCCCGTATA
AACCGCTGAACAAAACCGGCCTGGTGCTGAAAATGGATTTAGATCCGAACGATGTGACCTTTACCTATAGCCA
TTTTGCGGTGCCGCAGGCGAAAGGCAACAACGTGGTGATTACCAGCTATATGACCAACCGCGGCTTTTATGCG
GATAAACAGAGTACCTTTGCGCCGAGCTTTCTGCTGAACATTAAAGGCAAAAAAACCAGCGTGGTGAAAGATA
GCATTCTGGAACAGGGCCAACTTACCGTGAACAAAGGTACCTAATAGCTGAAGATCCGGCTGCTAACAAAGCC
CGAAAGGAAGCTGAGTTGGCTGCTGCCACCGCTGAGCAATAACTAGCATAACCCCTTGGGGCCTCTAAACGG
GTCTTGAGGGGTTTTTTGCTGAAAGGATCCACGTCAGGTTGCCATCGTATTAGCACCCCGCACTCTAGATTTCA
GTGCAATTTATCTCTTCAAATGTAGCACCTGAAGTCAGCCCCATACGATATAAGTTGTAATTCTCATGTTTGACA
GCTTATCATCGATAAGCTTTAATGCGGTAGTTTATCACAGTTAAATTGCTACCAATTTCTACTCTTGTAGATTGA
TCCCCGATCTGGAGAACACCAATTTCTACTCTTGTAGATAGCTGCCACATCAAACTGGAGAATAAGGTCAGTTT
CACCTGTTTTACGTAAAAACCCGCTTCGGCGGGTTTTTACTTTTGGCTCGAGCAGCTCGATGTTAGCGTCGGCG
TCCCGGGAGCTGCATGTGTCAGAGGTTTTCACCGTCATCACCGAAACGCGCGAGACGAAAGGGCCTCGTGATA
CGCCTATTTTTATAGGTTAATGTCATGATAATAATGGTTTCTTAGACGTCAGGTGGCACTTTTCGGGAAGTTCCT
ATTCTCTAGAAAGTATAGGAACTTCGGAAATGTGCGCGGAACCCCTATTTGTTTATTTTTCTAAATACATTCAAA
TATGTATCCGCTCATGAGACAATAACCCTGATAAATGCTTCAATAATATTGAAAAAGGAAGAGTATGATTGAAC
AAGATGGATTGCACGCAGGTTCTCCGGCCGCTTGGGTGGAGAGGCTATTCGGCTATGACTGGGCACAACAGA
CAATCGGCTGCTCTGATGCCGCCGTGTTCCGGCTGTCAGCGCAGGGGCGCCTGGTTCTTTTTGTCAAGACCGA
CCTCTCCGGTGCCCTGAATGAACTGCAAGACGAGGCAGCGCGGCTATCGTGGCTGGCCACGACGGGCGTTCCT
TGCGCAGCTGTGCTCGACGTTGTCACTGAAGCGGGAAGGGACTGGCTGCTATTGGGCGAAGTGCCGGGGCAG
GATCTCCTGTCATCTCACCTTGCTCCTGCCGAGAAAGTATCCATCATGGCTGATGCAATGCGGCGGCTGCATAC
GCTTGATCCGGCTACCTGCCCATTCGACCACCAAGCGAAACATCGCATCGAGCGAGCACGTACTCGGATGGAA
GCCGGTCTTGTCGATCAGGATGATCTGGACGAAGAGCATCAGGGGCTCGCGCCAGCCGAACTGTTCGCCAGG

CTCAAGGCGAGCATGCCCGACGGCGAGGATCTCGTCGTGACCCATGGCGATGCCTGCTTGCCGAATATCATGG
TGGAAAATGGCCGCTTTTCTGGATTCATCGACTGTGGCCGGCTGGGTGTGGCGGACCGCTATCAGGACATAGC
GTTGGCTACCCGTGATATTGCTGAAGAGCTTGGCGGCGAATGGGCTGACCGCTTCCTCGTGCTTTACGGTATC
GCCGCTCCCGATTCGCAGCGCATCGCCTTCTATCGCCTTCTTGACGAGTTCTTCTGACTGTCAGACCAAGTTTAC
TCATATATACTTTAGATTGATTTAAAACTTCATTTTTAATTTAAAAGGATCTAGGTGAAGATCCTTTTTGATAATC
TCATGACCAAAATCCCTTAACGTGAGTTTTCGTTCCACTGAAGTTCCTATTCTCTAGAAAGTATAGGAACTTCGA
GCGTCAGACCCCGTAGAAAAGATCAAAGGATCTTC

**>pGRNA-sacB-deoR**

TTGAGATCCTTTTTTTCTGTGCGTAATCTGCTGCTTGCAAACAAAAAAACCACCGCTACCAGCGGTGGTTTGTTT
GCCGGATCAAGAGCTACCAACTCTTTTTCCGAAGGTAACTGGCTTCAGCAGAGCGCAGATACCAAATACTGTTC
TTCTAGTGTAGCCGTAGTTAGGCCACCACTTCAAGAACTCTGTAGCACCGCCTACATACCTCGCTCTGCTAATCC
TGTTACCAGTGGCTGCTGCCAGTGGCGATAAGTCGTGTCTTACCGGGTTGGACTCAAGACGATAGTTACCGGA
TAAGGCGCAGCGGTCGGGCTGAACGGGGGGTTCGTGCACACAGCCCAGCTTGGAGCGAACGACCTACACCG
AACTGAGATACCTACAGCGTGAGCTATGAGAAAGCGCCACGCTTCCCGAAGGGAGAAAGGCGGACAGGTATC
CGGTAAGCGGCAGGGTCGGAACAGGAGAGCGCACGAGGGAGCTTCCAGGGGGAAACGCCTGGTATCTTTAT
AGTCCTGTCGGGTTTCGCCACCTCTGACTTGAGCGTCGATTTTTGTGATGCTCGTCAGGGGGGCGGAGCCTAT
GGAAAAACGCCAGCAACGCGGCCTTTTTACGGTTCCTGGCCTTTTGCTGGCCTTTTGCTCACATGTTCAAGCTTA
GAGGTTCCAACTTTCACCATAATGAAACACTCGAGAATAATTTTGTTTAACTTTAAGAAGGAGATATACATATG
AACATTAAAAAATTTGCGAAACAGGCGACAGTGCTGACCTTTACAACCGCGCTGTTAGCGGGCGGCGCGACCC
AGGCGTTTGCGAAAGAAACCAACCAGAAACCGTATAAAGAAACCTATGGCATTAGCCATATTACCCGCCATGA
TATGCTGCAGATCCCGGAACAGCAGAAAAACGAAAAATATCAGGTGCCGGAATTTGATAGCAGCACCATTAAA
AACATTAGCAGCGCGAAAGGCCTGGATGTGTGGGATAGCTGGCCGCTGCAGAACGCGGATGGCACCGTGGC
GAACTATCATGGCTATCATATTGTGTTTGCGCTGGCGGGCGATCCGAAAAACGCGGATGATACCAGTATTTAT
ATGTTTTATCAGAAAGTGGGCGAAACCAGCATTGATAGCTGGAAAAACGCGGGCCGCGTGTTTAAAGATAGC
GATAAATTTGATGCGAACGATAGCATTCTGAAAGATCAGACCCAGGAATGGTCAGGCAGCGCGACCTTTACCA
GCGATGGCAAAATTCGCCTGTTTTATACCGATTTTAGCGGCAAACATTATGGCAAACAGACCCTGACCACCGCC
CAGGTGAACGTGAGCGCGAGCGATAGCAGCCTGAACATTAACGGCGTGGAGGACTACAAGAGCATTTTTGAT
GGCGATGGCAAAACCTATCAGAACGTGCAACAGTTTATTGATGAAGGCAACTATAGCAGCGGCGATAACCATA
CCCTGCGCGATCCGCATTATGTGGAAGATAAAGGCCATAAATATCTGGTGTTTGAAGCGAACACCGGCACCGA
AGATGGCTATCAGGGCGAAGAAAGCCTGTTTAACAAAGCGTATTATGGCAAAAGCACCAGCTTTTTTCGCCAG
GAAAGCCAGAAACTGCTGCAGAGCGATAAAAAACGCACCGCGGAACTGGCGAACGGTGCGCTGGGCATGAT
TGAACTGAACGATGATTATACCCTGAAAAAAGTGATGAAACCGCTGATTGCGAGCAACACAGTGACAGATGA
AATTGAACGCGCGAATGTTTTTAAAATGAACGGCAAATGGTATCTGTTTACCGATAGCCGCGGCAGCAAAATG
ACCATTGATGGCATTACCAGCAACGATATTTATATGCTGGGCTATGTGAGCAACAGTCTGACAGGCCCGTATA
AACCGCTGAACAAAACCGGCCTGGTGCTGAAAATGGATTTAGATCCGAACGATGTGACCTTTACCTATAGCCA
TTTTGCGGTGCCGCAGGCGAAAGGCAACAACGTGGTGATTACCAGCTATATGACCAACCGCGGCTTTTATGCG
GATAAACAGAGTACCTTTGCGCCGAGCTTTCTGCTGAACATTAAAGGCAAAAAAACCAGCGTGGTGAAAGATA
GCATTCTGGAACAGGGCCAACTTACCGTGAACAAAGGTACCTAATAGCTGAAGATCCGGCTGCTAACAAAGCC
CGAAAGGAAGCTGAGTTGGCTGCTGCCACCGCTGAGCAATAACTAGCATAACCCCTTGGGGCCTCTAAACGG
GTCTTGAGGGGTTTTTTGCTGAAAGGATCCACGTCAGGTTGCCATCGTATTAGCACCCCGCACTCTAGATTTCA
GTGCAATTTATCTCTTCAAATGTAGCACCTGAAGTCAGCCCCATACGATATAAGTTGTAATTCTCATGTTTGACA
GCTTATCATCGATAAGCTTTAATGCGGTAGTTTATCACAGTTAAATTGCTACCAATTTCTACTCTTGTAGATAAG
AGTTGCCGGTAAAACACTGGAATTTCTACTCTTGTAGATAGCTGCCACATCAAACTGGAGAATAAGGTCAGTTT
CACCTGTTTTACGTAAAAACCCGCTTCGGCGGGTTTTTACTTTTGGCTCGAGCAGCTCGATGTTAGCGTCGGCG
TCCCGGGAGCTGCATGTGTCAGAGGTTTTCACCGTCATCACCGAAACGCGCGAGACGAAAGGGCCTCGTGATA
CGCCTATTTTTATAGGTTAATGTCATGATAATAATGGTTTCTTAGACGTCAGGTGGCACTTTTCGGGAAGTTCCT
ATTCTCTAGAAAGTATAGGAACTTCGGAAATGTGCGCGGAACCCCTATTTGTTTATTTTTCTAAATACATTCAAA
TATGTATCCGCTCATGAGACAATAACCCTGATAAATGCTTCAATAATATTGAAAAAGGAAGAGTATGATTGAAC
AAGATGGATTGCACGCAGGTTCTCCGGCCGCTTGGGTGGAGAGGCTATTCGGCTATGACTGGGCACAACAGA
CAATCGGCTGCTCTGATGCCGCCGTGTTCCGGCTGTCAGCGCAGGGGCGCCTGGTTCTTTTTGTCAAGACCGA
CCTCTCCGGTGCCCTGAATGAACTGCAAGACGAGGCAGCGCGGCTATCGTGGCTGGCCACGACGGGCGTTCCT
TGCGCAGCTGTGCTCGACGTTGTCACTGAAGCGGGAAGGGACTGGCTGCTATTGGGCGAAGTGCCGGGGCAG
GATCTCCTGTCATCTCACCTTGCTCCTGCCGAGAAAGTATCCATCATGGCTGATGCAATGCGGCGGCTGCATAC
GCTTGATCCGGCTACCTGCCCATTCGACCACCAAGCGAAACATCGCATCGAGCGAGCACGTACTCGGATGGAA
GCCGGTCTTGTCGATCAGGATGATCTGGACGAAGAGCATCAGGGGCTCGCGCCAGCCGAACTGTTCGCCAGG

CTCAAGGCGAGCATGCCCGACGGCGAGGATCTCGTCGTGACCCATGGCGATGCCTGCTTGCCGAATATCATGG
TGGAAAATGGCCGCTTTTCTGGATTCATCGACTGTGGCCGGCTGGGTGTGGCGGACCGCTATCAGGACATAGC
GTTGGCTACCCGTGATATTGCTGAAGAGCTTGGCGGCGAATGGGCTGACCGCTTCCTCGTGCTTTACGGTATC
GCCGCTCCCGATTCGCAGCGCATCGCCTTCTATCGCCTTCTTGACGAGTTCTTCTGACTGTCAGACCAAGTTTAC
TCATATATACTTTAGATTGATTTAAAACTTCATTTTTAATTTAAAAGGATCTAGGTGAAGATCCTTTTTGATAATC
TCATGACCAAAATCCCTTAACGTGAGTTTTCGTTCCACTGAAGTTCCTATTCTCTAGAAAGTATAGGAACTTCGA
GCGTCAGACCCCGTAGAAAAGATCAAAGGATCTTC

**>pGRNA-sacB-relA**

TTGAGATCCTTTTTTTCTGTGCGTAATCTGCTGCTTGCAAACAAAAAAACCACCGCTACCAGCGGTGGTTTGTTT
GCCGGATCAAGAGCTACCAACTCTTTTTCCGAAGGTAACTGGCTTCAGCAGAGCGCAGATACCAAATACTGTTC
TTCTAGTGTAGCCGTAGTTAGGCCACCACTTCAAGAACTCTGTAGCACCGCCTACATACCTCGCTCTGCTAATCC
TGTTACCAGTGGCTGCTGCCAGTGGCGATAAGTCGTGTCTTACCGGGTTGGACTCAAGACGATAGTTACCGGA
TAAGGCGCAGCGGTCGGGCTGAACGGGGGGTTCGTGCACACAGCCCAGCTTGGAGCGAACGACCTACACCG
AACTGAGATACCTACAGCGTGAGCTATGAGAAAGCGCCACGCTTCCCGAAGGGAGAAAGGCGGACAGGTATC
CGGTAAGCGGCAGGGTCGGAACAGGAGAGCGCACGAGGGAGCTTCCAGGGGGAAACGCCTGGTATCTTTAT
AGTCCTGTCGGGTTTCGCCACCTCTGACTTGAGCGTCGATTTTTGTGATGCTCGTCAGGGGGGCGGAGCCTAT
GGAAAAACGCCAGCAACGCGGCCTTTTTACGGTTCCTGGCCTTTTGCTGGCCTTTTGCTCACATGTTCAAGCTTA
GAGGTTCCAACTTTCACCATAATGAAACACTCGAGAATAATTTTGTTTAACTTTAAGAAGGAGATATACATATG
AACATTAAAAAATTTGCGAAACAGGCGACAGTGCTGACCTTTACAACCGCGCTGTTAGCGGGCGGCGCGACCC
AGGCGTTTGCGAAAGAAACCAACCAGAAACCGTATAAAGAAACCTATGGCATTAGCCATATTACCCGCCATGA
TATGCTGCAGATCCCGGAACAGCAGAAAAACGAAAAATATCAGGTGCCGGAATTTGATAGCAGCACCATTAAA
AACATTAGCAGCGCGAAAGGCCTGGATGTGTGGGATAGCTGGCCGCTGCAGAACGCGGATGGCACCGTGGC
GAACTATCATGGCTATCATATTGTGTTTGCGCTGGCGGGCGATCCGAAAAACGCGGATGATACCAGTATTTAT
ATGTTTTATCAGAAAGTGGGCGAAACCAGCATTGATAGCTGGAAAAACGCGGGCCGCGTGTTTAAAGATAGC
GATAAATTTGATGCGAACGATAGCATTCTGAAAGATCAGACCCAGGAATGGTCAGGCAGCGCGACCTTTACCA
GCGATGGCAAAATTCGCCTGTTTTATACCGATTTTAGCGGCAAACATTATGGCAAACAGACCCTGACCACCGCC
CAGGTGAACGTGAGCGCGAGCGATAGCAGCCTGAACATTAACGGCGTGGAGGACTACAAGAGCATTTTTGAT
GGCGATGGCAAAACCTATCAGAACGTGCAACAGTTTATTGATGAAGGCAACTATAGCAGCGGCGATAACCATA
CCCTGCGCGATCCGCATTATGTGGAAGATAAAGGCCATAAATATCTGGTGTTTGAAGCGAACACCGGCACCGA
AGATGGCTATCAGGGCGAAGAAAGCCTGTTTAACAAAGCGTATTATGGCAAAAGCACCAGCTTTTTTCGCCAG
GAAAGCCAGAAACTGCTGCAGAGCGATAAAAAACGCACCGCGGAACTGGCGAACGGTGCGCTGGGCATGAT
TGAACTGAACGATGATTATACCCTGAAAAAAGTGATGAAACCGCTGATTGCGAGCAACACAGTGACAGATGA
AATTGAACGCGCGAATGTTTTTAAAATGAACGGCAAATGGTATCTGTTTACCGATAGCCGCGGCAGCAAAATG
ACCATTGATGGCATTACCAGCAACGATATTTATATGCTGGGCTATGTGAGCAACAGTCTGACAGGCCCGTATA
AACCGCTGAACAAAACCGGCCTGGTGCTGAAAATGGATTTAGATCCGAACGATGTGACCTTTACCTATAGCCA
TTTTGCGGTGCCGCAGGCGAAAGGCAACAACGTGGTGATTACCAGCTATATGACCAACCGCGGCTTTTATGCG
GATAAACAGAGTACCTTTGCGCCGAGCTTTCTGCTGAACATTAAAGGCAAAAAAACCAGCGTGGTGAAAGATA
GCATTCTGGAACAGGGCCAACTTACCGTGAACAAAGGTACCTAATAGCTGAAGATCCGGCTGCTAACAAAGCC
CGAAAGGAAGCTGAGTTGGCTGCTGCCACCGCTGAGCAATAACTAGCATAACCCCTTGGGGCCTCTAAACGG
GTCTTGAGGGGTTTTTTGCTGAAAGGATCCACGTCAGGTTGCCATCGTATTAGCACCCCGCACTCTAGATTTCA
GTGCAATTTATCTCTTCAAATGTAGCACCTGAAGTCAGCCCCATACGATATAAGTTGTAATTCTCATGTTTGACA
GCTTATCATCGATAAGCTTTAATGCGGTAGTTTATCACAGTTAAATTGCTACCAATTTCTACTCTTGTAGATATTA
CCCAGGGGCGCGGTATTTCAATTTCTACTCTTGTAGATAGCTGCCACATCAAACTGGAGAATAAGGTCAGTTTC
ACCTGTTTTACGTAAAAACCCGCTTCGGCGGGTTTTTACTTTTGGCTCGAGCAGCTCGATGTTAGCGTCGGCGT
CCCGGGAGCTGCATGTGTCAGAGGTTTTCACCGTCATCACCGAAACGCGCGAGACGAAAGGGCCTCGTGATA
CGCCTATTTTTATAGGTTAATGTCATGATAATAATGGTTTCTTAGACGTCAGGTGGCACTTTTCGGGAAGTTCCT
ATTCTCTAGAAAGTATAGGAACTTCGGAAATGTGCGCGGAACCCCTATTTGTTTATTTTTCTAAATACATTCAAA
TATGTATCCGCTCATGAGACAATAACCCTGATAAATGCTTCAATAATATTGAAAAAGGAAGAGTATGATTGAAC
AAGATGGATTGCACGCAGGTTCTCCGGCCGCTTGGGTGGAGAGGCTATTCGGCTATGACTGGGCACAACAGA
CAATCGGCTGCTCTGATGCCGCCGTGTTCCGGCTGTCAGCGCAGGGGCGCCTGGTTCTTTTTGTCAAGACCGA
CCTCTCCGGTGCCCTGAATGAACTGCAAGACGAGGCAGCGCGGCTATCGTGGCTGGCCACGACGGGCGTTCCT
TGCGCAGCTGTGCTCGACGTTGTCACTGAAGCGGGAAGGGACTGGCTGCTATTGGGCGAAGTGCCGGGGCAG
GATCTCCTGTCATCTCACCTTGCTCCTGCCGAGAAAGTATCCATCATGGCTGATGCAATGCGGCGGCTGCATAC
GCTTGATCCGGCTACCTGCCCATTCGACCACCAAGCGAAACATCGCATCGAGCGAGCACGTACTCGGATGGAA
GCCGGTCTTGTCGATCAGGATGATCTGGACGAAGAGCATCAGGGGCTCGCGCCAGCCGAACTGTTCGCCAGG

CTCAAGGCGAGCATGCCCGACGGCGAGGATCTCGTCGTGACCCATGGCGATGCCTGCTTGCCGAATATCATGG
TGGAAAATGGCCGCTTTTCTGGATTCATCGACTGTGGCCGGCTGGGTGTGGCGGACCGCTATCAGGACATAGC
GTTGGCTACCCGTGATATTGCTGAAGAGCTTGGCGGCGAATGGGCTGACCGCTTCCTCGTGCTTTACGGTATC
GCCGCTCCCGATTCGCAGCGCATCGCCTTCTATCGCCTTCTTGACGAGTTCTTCTGACTGTCAGACCAAGTTTAC
TCATATATACTTTAGATTGATTTAAAACTTCATTTTTAATTTAAAAGGATCTAGGTGAAGATCCTTTTTGATAATC
TCATGACCAAAATCCCTTAACGTGAGTTTTCGTTCCACTGAAGTTCCTATTCTCTAGAAAGTATAGGAACTTCGA
GCGTCAGACCCCGTAGAAAAGATCAAAGGATCTTC

**>pGRNA-sacB-nupG**

TTGAGATCCTTTTTTTCTGTGCGTAATCTGCTGCTTGCAAACAAAAAAACCACCGCTACCAGCGGTGGTTTGTTT
GCCGGATCAAGAGCTACCAACTCTTTTTCCGAAGGTAACTGGCTTCAGCAGAGCGCAGATACCAAATACTGTTC
TTCTAGTGTAGCCGTAGTTAGGCCACCACTTCAAGAACTCTGTAGCACCGCCTACATACCTCGCTCTGCTAATCC
TGTTACCAGTGGCTGCTGCCAGTGGCGATAAGTCGTGTCTTACCGGGTTGGACTCAAGACGATAGTTACCGGA
TAAGGCGCAGCGGTCGGGCTGAACGGGGGGTTCGTGCACACAGCCCAGCTTGGAGCGAACGACCTACACCG
AACTGAGATACCTACAGCGTGAGCTATGAGAAAGCGCCACGCTTCCCGAAGGGAGAAAGGCGGACAGGTATC
CGGTAAGCGGCAGGGTCGGAACAGGAGAGCGCACGAGGGAGCTTCCAGGGGGAAACGCCTGGTATCTTTAT
AGTCCTGTCGGGTTTCGCCACCTCTGACTTGAGCGTCGATTTTTGTGATGCTCGTCAGGGGGGCGGAGCCTAT
GGAAAAACGCCAGCAACGCGGCCTTTTTACGGTTCCTGGCCTTTTGCTGGCCTTTTGCTCACATGTTCAAGCTTA
GAGGTTCCAACTTTCACCATAATGAAACACTCGAGAATAATTTTGTTTAACTTTAAGAAGGAGATATACATATG
AACATTAAAAAATTTGCGAAACAGGCGACAGTGCTGACCTTTACAACCGCGCTGTTAGCGGGCGGCGCGACCC
AGGCGTTTGCGAAAGAAACCAACCAGAAACCGTATAAAGAAACCTATGGCATTAGCCATATTACCCGCCATGA
TATGCTGCAGATCCCGGAACAGCAGAAAAACGAAAAATATCAGGTGCCGGAATTTGATAGCAGCACCATTAAA
AACATTAGCAGCGCGAAAGGCCTGGATGTGTGGGATAGCTGGCCGCTGCAGAACGCGGATGGCACCGTGGC
GAACTATCATGGCTATCATATTGTGTTTGCGCTGGCGGGCGATCCGAAAAACGCGGATGATACCAGTATTTAT
ATGTTTTATCAGAAAGTGGGCGAAACCAGCATTGATAGCTGGAAAAACGCGGGCCGCGTGTTTAAAGATAGC
GATAAATTTGATGCGAACGATAGCATTCTGAAAGATCAGACCCAGGAATGGTCAGGCAGCGCGACCTTTACCA
GCGATGGCAAAATTCGCCTGTTTTATACCGATTTTAGCGGCAAACATTATGGCAAACAGACCCTGACCACCGCC
CAGGTGAACGTGAGCGCGAGCGATAGCAGCCTGAACATTAACGGCGTGGAGGACTACAAGAGCATTTTTGAT
GGCGATGGCAAAACCTATCAGAACGTGCAACAGTTTATTGATGAAGGCAACTATAGCAGCGGCGATAACCATA
CCCTGCGCGATCCGCATTATGTGGAAGATAAAGGCCATAAATATCTGGTGTTTGAAGCGAACACCGGCACCGA
AGATGGCTATCAGGGCGAAGAAAGCCTGTTTAACAAAGCGTATTATGGCAAAAGCACCAGCTTTTTTCGCCAG
GAAAGCCAGAAACTGCTGCAGAGCGATAAAAAACGCACCGCGGAACTGGCGAACGGTGCGCTGGGCATGAT
TGAACTGAACGATGATTATACCCTGAAAAAAGTGATGAAACCGCTGATTGCGAGCAACACAGTGACAGATGA
AATTGAACGCGCGAATGTTTTTAAAATGAACGGCAAATGGTATCTGTTTACCGATAGCCGCGGCAGCAAAATG
ACCATTGATGGCATTACCAGCAACGATATTTATATGCTGGGCTATGTGAGCAACAGTCTGACAGGCCCGTATA
AACCGCTGAACAAAACCGGCCTGGTGCTGAAAATGGATTTAGATCCGAACGATGTGACCTTTACCTATAGCCA
TTTTGCGGTGCCGCAGGCGAAAGGCAACAACGTGGTGATTACCAGCTATATGACCAACCGCGGCTTTTATGCG
GATAAACAGAGTACCTTTGCGCCGAGCTTTCTGCTGAACATTAAAGGCAAAAAAACCAGCGTGGTGAAAGATA
GCATTCTGGAACAGGGCCAACTTACCGTGAACAAAGGTACCTAATAGCTGAAGATCCGGCTGCTAACAAAGCC
CGAAAGGAAGCTGAGTTGGCTGCTGCCACCGCTGAGCAATAACTAGCATAACCCCTTGGGGCCTCTAAACGG
GTCTTGAGGGGTTTTTTGCTGAAAGGATCCACGTCAGGTTGCCATCGTATTAGCACCCCGCACTCTAGATTTCA
GTGCAATTTATCTCTTCAAATGTAGCACCTGAAGTCAGCCCCATACGATATAAGTTGTAATTCTCATGTTTGACA
GCTTATCATCGATAAGCTTTAATGCGGTAGTTTATCACAGTTAAATTGCTACCAATTTCTACTCTTGTAGATTAG
CTCACTGGGTATCGCAGCGGAATTTCTACTCTTGTAGATAGCTGCCACATCAAACTGGAGAATAAGGTCAGTTT
CACCTGTTTTACGTAAAAACCCGCTTCGGCGGGTTTTTACTTTTGGCTCGAGCAGCTCGATGTTAGCGTCGGCG
TCCCGGGAGCTGCATGTGTCAGAGGTTTTCACCGTCATCACCGAAACGCGCGAGACGAAAGGGCCTCGTGATA
CGCCTATTTTTATAGGTTAATGTCATGATAATAATGGTTTCTTAGACGTCAGGTGGCACTTTTCGGGAAGTTCCT
ATTCTCTAGAAAGTATAGGAACTTCGGAAATGTGCGCGGAACCCCTATTTGTTTATTTTTCTAAATACATTCAAA
TATGTATCCGCTCATGAGACAATAACCCTGATAAATGCTTCAATAATATTGAAAAAGGAAGAGTATGATTGAAC
AAGATGGATTGCACGCAGGTTCTCCGGCCGCTTGGGTGGAGAGGCTATTCGGCTATGACTGGGCACAACAGA
CAATCGGCTGCTCTGATGCCGCCGTGTTCCGGCTGTCAGCGCAGGGGCGCCTGGTTCTTTTTGTCAAGACCGA
CCTCTCCGGTGCCCTGAATGAACTGCAAGACGAGGCAGCGCGGCTATCGTGGCTGGCCACGACGGGCGTTCCT
TGCGCAGCTGTGCTCGACGTTGTCACTGAAGCGGGAAGGGACTGGCTGCTATTGGGCGAAGTGCCGGGGCAG
GATCTCCTGTCATCTCACCTTGCTCCTGCCGAGAAAGTATCCATCATGGCTGATGCAATGCGGCGGCTGCATAC
GCTTGATCCGGCTACCTGCCCATTCGACCACCAAGCGAAACATCGCATCGAGCGAGCACGTACTCGGATGGAA
GCCGGTCTTGTCGATCAGGATGATCTGGACGAAGAGCATCAGGGGCTCGCGCCAGCCGAACTGTTCGCCAGG

CTCAAGGCGAGCATGCCCGACGGCGAGGATCTCGTCGTGACCCATGGCGATGCCTGCTTGCCGAATATCATGG
TGGAAAATGGCCGCTTTTCTGGATTCATCGACTGTGGCCGGCTGGGTGTGGCGGACCGCTATCAGGACATAGC
GTTGGCTACCCGTGATATTGCTGAAGAGCTTGGCGGCGAATGGGCTGACCGCTTCCTCGTGCTTTACGGTATC
GCCGCTCCCGATTCGCAGCGCATCGCCTTCTATCGCCTTCTTGACGAGTTCTTCTGACTGTCAGACCAAGTTTAC
TCATATATACTTTAGATTGATTTAAAACTTCATTTTTAATTTAAAAGGATCTAGGTGAAGATCCTTTTTGATAATC
TCATGACCAAAATCCCTTAACGTGAGTTTTCGTTCCACTGAAGTTCCTATTCTCTAGAAAGTATAGGAACTTCGA
GCGTCAGACCCCGTAGAAAAGATCAAAGGATCTTC

**>pGRNA-sacB-PgalP**

TTGAGATCCTTTTTTTCTGTGCGTAATCTGCTGCTTGCAAACAAAAAAACCACCGCTACCAGCGGTGGTTTGTTT
GCCGGATCAAGAGCTACCAACTCTTTTTCCGAAGGTAACTGGCTTCAGCAGAGCGCAGATACCAAATACTGTTC
TTCTAGTGTAGCCGTAGTTAGGCCACCACTTCAAGAACTCTGTAGCACCGCCTACATACCTCGCTCTGCTAATCC
TGTTACCAGTGGCTGCTGCCAGTGGCGATAAGTCGTGTCTTACCGGGTTGGACTCAAGACGATAGTTACCGGA
TAAGGCGCAGCGGTCGGGCTGAACGGGGGGTTCGTGCACACAGCCCAGCTTGGAGCGAACGACCTACACCG
AACTGAGATACCTACAGCGTGAGCTATGAGAAAGCGCCACGCTTCCCGAAGGGAGAAAGGCGGACAGGTATC
CGGTAAGCGGCAGGGTCGGAACAGGAGAGCGCACGAGGGAGCTTCCAGGGGGAAACGCCTGGTATCTTTAT
AGTCCTGTCGGGTTTCGCCACCTCTGACTTGAGCGTCGATTTTTGTGATGCTCGTCAGGGGGGCGGAGCCTAT
GGAAAAACGCCAGCAACGCGGCCTTTTTACGGTTCCTGGCCTTTTGCTGGCCTTTTGCTCACATGTTCAAGCTTA
GAGGTTCCAACTTTCACCATAATGAAACACTCGAGAATAATTTTGTTTAACTTTAAGAAGGAGATATACATATG
AACATTAAAAAATTTGCGAAACAGGCGACAGTGCTGACCTTTACAACCGCGCTGTTAGCGGGCGGCGCGACCC
AGGCGTTTGCGAAAGAAACCAACCAGAAACCGTATAAAGAAACCTATGGCATTAGCCATATTACCCGCCATGA
TATGCTGCAGATCCCGGAACAGCAGAAAAACGAAAAATATCAGGTGCCGGAATTTGATAGCAGCACCATTAAA
AACATTAGCAGCGCGAAAGGCCTGGATGTGTGGGATAGCTGGCCGCTGCAGAACGCGGATGGCACCGTGGC
GAACTATCATGGCTATCATATTGTGTTTGCGCTGGCGGGCGATCCGAAAAACGCGGATGATACCAGTATTTAT
ATGTTTTATCAGAAAGTGGGCGAAACCAGCATTGATAGCTGGAAAAACGCGGGCCGCGTGTTTAAAGATAGC
GATAAATTTGATGCGAACGATAGCATTCTGAAAGATCAGACCCAGGAATGGTCAGGCAGCGCGACCTTTACCA
GCGATGGCAAAATTCGCCTGTTTTATACCGATTTTAGCGGCAAACATTATGGCAAACAGACCCTGACCACCGCC
CAGGTGAACGTGAGCGCGAGCGATAGCAGCCTGAACATTAACGGCGTGGAGGACTACAAGAGCATTTTTGAT
GGCGATGGCAAAACCTATCAGAACGTGCAACAGTTTATTGATGAAGGCAACTATAGCAGCGGCGATAACCATA
CCCTGCGCGATCCGCATTATGTGGAAGATAAAGGCCATAAATATCTGGTGTTTGAAGCGAACACCGGCACCGA
AGATGGCTATCAGGGCGAAGAAAGCCTGTTTAACAAAGCGTATTATGGCAAAAGCACCAGCTTTTTTCGCCAG
GAAAGCCAGAAACTGCTGCAGAGCGATAAAAAACGCACCGCGGAACTGGCGAACGGTGCGCTGGGCATGAT
TGAACTGAACGATGATTATACCCTGAAAAAAGTGATGAAACCGCTGATTGCGAGCAACACAGTGACAGATGA
AATTGAACGCGCGAATGTTTTTAAAATGAACGGCAAATGGTATCTGTTTACCGATAGCCGCGGCAGCAAAATG
ACCATTGATGGCATTACCAGCAACGATATTTATATGCTGGGCTATGTGAGCAACAGTCTGACAGGCCCGTATA
AACCGCTGAACAAAACCGGCCTGGTGCTGAAAATGGATTTAGATCCGAACGATGTGACCTTTACCTATAGCCA
TTTTGCGGTGCCGCAGGCGAAAGGCAACAACGTGGTGATTACCAGCTATATGACCAACCGCGGCTTTTATGCG
GATAAACAGAGTACCTTTGCGCCGAGCTTTCTGCTGAACATTAAAGGCAAAAAAACCAGCGTGGTGAAAGATA
GCATTCTGGAACAGGGCCAACTTACCGTGAACAAAGGTACCTAATAGCTGAAGATCCGGCTGCTAACAAAGCC
CGAAAGGAAGCTGAGTTGGCTGCTGCCACCGCTGAGCAATAACTAGCATAACCCCTTGGGGCCTCTAAACGG
GTCTTGAGGGGTTTTTTGCTGAAAGGATCCACGTCAGGTTGCCATCGTATTAGCACCCCGCACTCTAGATTTCA
GTGCAATTTATCTCTTCAAATGTAGCACCTGAAGTCAGCCCCATACGATATAAGTTGTAATTCTCATGTTTGACA
GCTTATCATCGATAAGCTTTAATGCGGTAGTTTATCACAGTTAAATTGCTACCAATTTCTACTCTTGTAGATTCTG
TTGTGGTTGGTGTAATCGGAATTTCTACTCTTGTAGATAGCTGCCACATCAAACTGGAGAATAAGGTCAGTTTC
ACCTGTTTTACGTAAAAACCCGCTTCGGCGGGTTTTTACTTTTGGCTCGAGCAGCTCGATGTTAGCGTCGGCGT
CCCGGGAGCTGCATGTGTCAGAGGTTTTCACCGTCATCACCGAAACGCGCGAGACGAAAGGGCCTCGTGATA
CGCCTATTTTTATAGGTTAATGTCATGATAATAATGGTTTCTTAGACGTCAGGTGGCACTTTTCGGGAAGTTCCT
ATTCTCTAGAAAGTATAGGAACTTCGGAAATGTGCGCGGAACCCCTATTTGTTTATTTTTCTAAATACATTCAAA
TATGTATCCGCTCATGAGACAATAACCCTGATAAATGCTTCAATAATATTGAAAAAGGAAGAGTATGATTGAAC
AAGATGGATTGCACGCAGGTTCTCCGGCCGCTTGGGTGGAGAGGCTATTCGGCTATGACTGGGCACAACAGA
CAATCGGCTGCTCTGATGCCGCCGTGTTCCGGCTGTCAGCGCAGGGGCGCCTGGTTCTTTTTGTCAAGACCGA
CCTCTCCGGTGCCCTGAATGAACTGCAAGACGAGGCAGCGCGGCTATCGTGGCTGGCCACGACGGGCGTTCCT
TGCGCAGCTGTGCTCGACGTTGTCACTGAAGCGGGAAGGGACTGGCTGCTATTGGGCGAAGTGCCGGGGCAG
GATCTCCTGTCATCTCACCTTGCTCCTGCCGAGAAAGTATCCATCATGGCTGATGCAATGCGGCGGCTGCATAC
GCTTGATCCGGCTACCTGCCCATTCGACCACCAAGCGAAACATCGCATCGAGCGAGCACGTACTCGGATGGAA
GCCGGTCTTGTCGATCAGGATGATCTGGACGAAGAGCATCAGGGGCTCGCGCCAGCCGAACTGTTCGCCAGG

CTCAAGGCGAGCATGCCCGACGGCGAGGATCTCGTCGTGACCCATGGCGATGCCTGCTTGCCGAATATCATGG
TGGAAAATGGCCGCTTTTCTGGATTCATCGACTGTGGCCGGCTGGGTGTGGCGGACCGCTATCAGGACATAGC
GTTGGCTACCCGTGATATTGCTGAAGAGCTTGGCGGCGAATGGGCTGACCGCTTCCTCGTGCTTTACGGTATC
GCCGCTCCCGATTCGCAGCGCATCGCCTTCTATCGCCTTCTTGACGAGTTCTTCTGACTGTCAGACCAAGTTTAC
TCATATATACTTTAGATTGATTTAAAACTTCATTTTTAATTTAAAAGGATCTAGGTGAAGATCCTTTTTGATAATC
TCATGACCAAAATCCCTTAACGTGAGTTTTCGTTCCACTGAAGTTCCTATTCTCTAGAAAGTATAGGAACTTCGA
GCGTCAGACCCCGTAGAAAAGATCAAAGGATCTTC

**>pGRNA-sacB-ptsH**

TTGAGATCCTTTTTTTCTGTGCGTAATCTGCTGCTTGCAAACAAAAAAACCACCGCTACCAGCGGTGGTTTGTTT
GCCGGATCAAGAGCTACCAACTCTTTTTCCGAAGGTAACTGGCTTCAGCAGAGCGCAGATACCAAATACTGTTC
TTCTAGTGTAGCCGTAGTTAGGCCACCACTTCAAGAACTCTGTAGCACCGCCTACATACCTCGCTCTGCTAATCC
TGTTACCAGTGGCTGCTGCCAGTGGCGATAAGTCGTGTCTTACCGGGTTGGACTCAAGACGATAGTTACCGGA
TAAGGCGCAGCGGTCGGGCTGAACGGGGGGTTCGTGCACACAGCCCAGCTTGGAGCGAACGACCTACACCG
AACTGAGATACCTACAGCGTGAGCTATGAGAAAGCGCCACGCTTCCCGAAGGGAGAAAGGCGGACAGGTATC
CGGTAAGCGGCAGGGTCGGAACAGGAGAGCGCACGAGGGAGCTTCCAGGGGGAAACGCCTGGTATCTTTAT
AGTCCTGTCGGGTTTCGCCACCTCTGACTTGAGCGTCGATTTTTGTGATGCTCGTCAGGGGGGCGGAGCCTAT
GGAAAAACGCCAGCAACGCGGCCTTTTTACGGTTCCTGGCCTTTTGCTGGCCTTTTGCTCACATGTTCAAGCTTA
GAGGTTCCAACTTTCACCATAATGAAACACTCGAGAATAATTTTGTTTAACTTTAAGAAGGAGATATACATATG
AACATTAAAAAATTTGCGAAACAGGCGACAGTGCTGACCTTTACAACCGCGCTGTTAGCGGGCGGCGCGACCC
AGGCGTTTGCGAAAGAAACCAACCAGAAACCGTATAAAGAAACCTATGGCATTAGCCATATTACCCGCCATGA
TATGCTGCAGATCCCGGAACAGCAGAAAAACGAAAAATATCAGGTGCCGGAATTTGATAGCAGCACCATTAAA
AACATTAGCAGCGCGAAAGGCCTGGATGTGTGGGATAGCTGGCCGCTGCAGAACGCGGATGGCACCGTGGC
GAACTATCATGGCTATCATATTGTGTTTGCGCTGGCGGGCGATCCGAAAAACGCGGATGATACCAGTATTTAT
ATGTTTTATCAGAAAGTGGGCGAAACCAGCATTGATAGCTGGAAAAACGCGGGCCGCGTGTTTAAAGATAGC
GATAAATTTGATGCGAACGATAGCATTCTGAAAGATCAGACCCAGGAATGGTCAGGCAGCGCGACCTTTACCA
GCGATGGCAAAATTCGCCTGTTTTATACCGATTTTAGCGGCAAACATTATGGCAAACAGACCCTGACCACCGCC
CAGGTGAACGTGAGCGCGAGCGATAGCAGCCTGAACATTAACGGCGTGGAGGACTACAAGAGCATTTTTGAT
GGCGATGGCAAAACCTATCAGAACGTGCAACAGTTTATTGATGAAGGCAACTATAGCAGCGGCGATAACCATA
CCCTGCGCGATCCGCATTATGTGGAAGATAAAGGCCATAAATATCTGGTGTTTGAAGCGAACACCGGCACCGA
AGATGGCTATCAGGGCGAAGAAAGCCTGTTTAACAAAGCGTATTATGGCAAAAGCACCAGCTTTTTTCGCCAG
GAAAGCCAGAAACTGCTGCAGAGCGATAAAAAACGCACCGCGGAACTGGCGAACGGTGCGCTGGGCATGAT
TGAACTGAACGATGATTATACCCTGAAAAAAGTGATGAAACCGCTGATTGCGAGCAACACAGTGACAGATGA
AATTGAACGCGCGAATGTTTTTAAAATGAACGGCAAATGGTATCTGTTTACCGATAGCCGCGGCAGCAAAATG
ACCATTGATGGCATTACCAGCAACGATATTTATATGCTGGGCTATGTGAGCAACAGTCTGACAGGCCCGTATA
AACCGCTGAACAAAACCGGCCTGGTGCTGAAAATGGATTTAGATCCGAACGATGTGACCTTTACCTATAGCCA
TTTTGCGGTGCCGCAGGCGAAAGGCAACAACGTGGTGATTACCAGCTATATGACCAACCGCGGCTTTTATGCG
GATAAACAGAGTACCTTTGCGCCGAGCTTTCTGCTGAACATTAAAGGCAAAAAAACCAGCGTGGTGAAAGATA
GCATTCTGGAACAGGGCCAACTTACCGTGAACAAAGGTACCTAATAGCTGAAGATCCGGCTGCTAACAAAGCC
CGAAAGGAAGCTGAGTTGGCTGCTGCCACCGCTGAGCAATAACTAGCATAACCCCTTGGGGCCTCTAAACGG
GTCTTGAGGGGTTTTTTGCTGAAAGGATCCACGTCAGGTTGCCATCGTATTAGCACCCCGCACTCTAGATTTCA
GTGCAATTTATCTCTTCAAATGTAGCACCTGAAGTCAGCCCCATACGATATAAGTTGTAATTCTCATGTTTGACA
GCTTATCATCGATAAGCTTTAATGCGGTAGTTTATCACAGTTAAATTGCTACCAATTTCTACTCTTGTAGATAAA
CTTTCGCCCCTCCTGGCATTAATTTCTACTCTTGTAGATAGCTGCCACATCAAACTGGAGAATAAGGTCAGTTTC
ACCTGTTTTACGTAAAAACCCGCTTCGGCGGGTTTTTACTTTTGGCTCGAGCAGCTCGATGTTAGCGTCGGCGT
CCCGGGAGCTGCATGTGTCAGAGGTTTTCACCGTCATCACCGAAACGCGCGAGACGAAAGGGCCTCGTGATA
CGCCTATTTTTATAGGTTAATGTCATGATAATAATGGTTTCTTAGACGTCAGGTGGCACTTTTCGGGAAGTTCCT
ATTCTCTAGAAAGTATAGGAACTTCGGAAATGTGCGCGGAACCCCTATTTGTTTATTTTTCTAAATACATTCAAA
TATGTATCCGCTCATGAGACAATAACCCTGATAAATGCTTCAATAATATTGAAAAAGGAAGAGTATGATTGAAC
AAGATGGATTGCACGCAGGTTCTCCGGCCGCTTGGGTGGAGAGGCTATTCGGCTATGACTGGGCACAACAGA
CAATCGGCTGCTCTGATGCCGCCGTGTTCCGGCTGTCAGCGCAGGGGCGCCTGGTTCTTTTTGTCAAGACCGA
CCTCTCCGGTGCCCTGAATGAACTGCAAGACGAGGCAGCGCGGCTATCGTGGCTGGCCACGACGGGCGTTCCT
TGCGCAGCTGTGCTCGACGTTGTCACTGAAGCGGGAAGGGACTGGCTGCTATTGGGCGAAGTGCCGGGGCAG
GATCTCCTGTCATCTCACCTTGCTCCTGCCGAGAAAGTATCCATCATGGCTGATGCAATGCGGCGGCTGCATAC
GCTTGATCCGGCTACCTGCCCATTCGACCACCAAGCGAAACATCGCATCGAGCGAGCACGTACTCGGATGGAA
GCCGGTCTTGTCGATCAGGATGATCTGGACGAAGAGCATCAGGGGCTCGCGCCAGCCGAACTGTTCGCCAGG

CTCAAGGCGAGCATGCCCGACGGCGAGGATCTCGTCGTGACCCATGGCGATGCCTGCTTGCCGAATATCATGG
TGGAAAATGGCCGCTTTTCTGGATTCATCGACTGTGGCCGGCTGGGTGTGGCGGACCGCTATCAGGACATAGC
GTTGGCTACCCGTGATATTGCTGAAGAGCTTGGCGGCGAATGGGCTGACCGCTTCCTCGTGCTTTACGGTATC
GCCGCTCCCGATTCGCAGCGCATCGCCTTCTATCGCCTTCTTGACGAGTTCTTCTGACTGTCAGACCAAGTTTAC
TCATATATACTTTAGATTGATTTAAAACTTCATTTTTAATTTAAAAGGATCTAGGTGAAGATCCTTTTTGATAATC
TCATGACCAAAATCCCTTAACGTGAGTTTTCGTTCCACTGAAGTTCCTATTCTCTAGAAAGTATAGGAACTTCGA
GCGTCAGACCCCGTAGAAAAGATCAAAGGATCTTC
